# Evolutionary selection of DNA nanostructures for cellular uptake

**DOI:** 10.64898/2026.02.20.706810

**Authors:** Anjali Rajwar, Lisa Eichhorn, Jakub Palacka, Sandra Ly, Erik Benson

**Affiliations:** Science for Life Laboratory (SciLifeLab), Department of Microbiology, Tumor and Cell Biology, Karolinska Institutet, Solna, Sweden

**Keywords:** DNA nanotechnology, cellular uptake, selection, DNA library

## Abstract

DNA nanotechnology offers precise, biocompatible structures with strong potential for targeted drug delivery, yet current discovery approaches rely on testing individual designs, limiting exploration of structural diversity. Here, we introduce an evolutionary selection strategy that screens large libraries of DNA nanostructures, each folded from a single-stranded ‘structure genome’ compatible with amplification and sequencing. Cellular internalization is used as the selection pressure: libraries are incubated with mammalian cells, internalized structures are recovered from lysates, and the process is iterated across multiple rounds in HEK293T and RAW264.7 cells. High-throughput sequencing of recovered structure genomes reveals cell-type-specific enrichment patterns, enabling the identification of individual nanostructures with preferential uptake. Selected candidates were synthesized and evaluated as purified structures, confirming differential internalization by quantitative flow cytometry and microscopy. Turning DNA nanostructure discovery into a selection-based process, could enable high-throughput exploration of structural diversity and provide an alternative route to identify nanostructures with cell-specific uptake properties for biomedical applications.

## 1. Introduction

A key goal of nanomedicine is to develop structures and particles with structural, mechanical and functional properties allowing them to specifically interact with and be taken up by certain cells to deliver therapeutic payloads. DNA nanotechnology is a powerful method for producing self-assembled nanostructures from oligonucleotides with fine control of size, shape, mechanics as well offering the capacity to display patterns of other biomolecules such as proteins^1–6^. This high degree of precision has allowed for the study of how design choices such as size, geometry and mechanical properties control the interaction with cells, tissues and organisms. Previous studies have demonstrated the importance of structure–activity relationships in DNA nanostructure uptake. For example, Bastings et al. compared 11 distinct DNA origami designs, ranging from 50 to 400 nm in size, across multiple cell lines, and observed that larger, more compact structures were preferentially internalized relative to elongated, high–aspect-ratio particles. Uptake rates, however, were shown to depend more strongly on cell type than particle geometry, with phagocytic cells such as dendritic cells exhibiting prolonged uptake without reaching saturation over 12 hours.^7^

Using the modular DNA brick strategy, researchers further engineered eight distinct DNA nanoparticle designs, including rectangular and tubular geometries of varying dimensions, for systemic siRNA delivery. Although all nanoparticles exhibited enhanced internalization compared to naked DNA, a small rectangular construct 6H x 96BP-Rect (a 6-helix-wide, 96-base-pair-long rectangle) achieved the highest uptake efficiency, reinforcing the significance of size and shape in cellular internalization.^8^ Complementing these findings, another study reported that tetrahedral DNA nanostructures exhibited superior cellular internalization relative to other three-dimensional geometries tested across both two-dimensional and three-dimensional cell culture models. This enhanced uptake was further confirmed in more complex biological contexts, including tumor spheroid invasion assays and epithelial barrier crossing in zebrafish models.^9^ Together, multiple studies underscore the critical role of rational nanoparticle design, particularly shape and size, in optimizing cellular uptake for biomedical applications.^1,4,6,10–13^

Despite these advances, a key limitation in exploring DNA nanostructure design space lies in the reliance on testing individual nanostructures one at a time, which makes it difficult to explore the vast design space available when shape, size, and mechanics can all be tuned independently. Evolutionary selection strategies offer a potential solution by enabling high-throughput functional screening of large structural libraries within a single experimental context. Such approaches underlie the success of SELEX for aptamer discovery,^14–17^ as well as DNA-encoded chemical libraries for small-molecule drug discovery.^18–20^

However, unlike oligonucleotide-based aptamers, most DNA nanostructures are assembled from multiple component strands via tile-based^21^ or origami methods.^22^ When such assemblies are pooled, amplified, and sequenced, structural information is lost, and refolding libraries from amplified mixtures is unlikely, as component strands from different designs can misassemble. To overcome this, recent advances have explored rationally designed DNA^23^ or RNA^24^ nanostructures that fold from one strand, including constructs compatible with co-transcriptional folding. In these designs, distal regions of the strand are programmed to form secondary and tertiary interactions such as helices, junctions, and kissing loops. While linear DNA fragment libraries can be generated at scale, producing diverse DNA nanostructure libraries through this route remains challenging, as interacting domains must be both encoded within the same strand and spatially segregated to ensure correct folding. In this work, we introduce a new strategy for generating DNA nanostructure libraries based on the ligation of smaller pre-assembled nanostructure fragments of varying sizes and complexity. This approach yields highly diverse libraries in which each structure’s overall architecture is defined by the specific combination of fragments. The resulting structures are diverse in shape yet remain traceable by sequencing. We apply this library in cellular selection experiments to investigate whether internalization by mammalian cells occurs randomly across structures or if certain structural features are preferentially selected. By performing iterative rounds of selection in HEK293T and RAW264.7 cells, followed by nanopore sequencing of internalized structures, we examine enrichment dynamics and compare results with Illumina sequencing for cross-validation. Candidate structures identified from both cell types were further characterized to assess and compare their internalization efficiencies. We evaluated the cellular internalization of preferentially selected candidates across both selected cell lines as well as in the cell line A549 that was not used for selection. Candidates enriched through selection exhibited the highest levels of internalization in their respective cell lines, whereas randomly selected, non-enriched candidates showed comparatively greater uptake in A549 cells. Together, these findings demonstrate a scalable framework that not only rapidly identifies compact, ∼200–500 bp DNA nanostructures that are preferentially internalized but also establishes a generalizable pipeline for engineering DNA nanocarriers tailored to specific cell types, paving the way for systematic development of nanomedicines with programmable payload delivery and multifunctional therapeutic capabilities.

## 2. Results and Discussion

### 2.1 Design and optimization of a DNA nanostructure library

We developed a library of 36 DNA nanostructure fragments that varied in size and complexity, each designed to assemble from one to four synthetic oligonucleotides **(Table S1)**. The sequence of each fragment oligonucleotide was designed from a set of 20-nt sequences devoid of homopolymer stretches, balanced GC content, matching melting temperatures and optimized to avoid cross-interactions and secondary structure formation. The fragment set included three-way junctions, four-way junctions, hairpins including kissing loops of various sizes, and linear fragments of different lengths **(Figure 1A)**. For linear fragments, we varied the sizes ranging from 14 nt to 20 nt, while for bulged fragments we fixed the length at 20 nt and varied the bulge size from 1 to 5 nucleotides. In addition, an ‘amplification loop’ hairpin was designed to provide primer-binding sites for PCR and two embedded 10-nt unique molecular identifier (UMI) sequences for lineage tracing. All fragments were designed to feature AGT or ACT sticky-end overhangs to facilitate ligation. The ligation was designed to be directional: linear fragments featured one AGT and one ACT overhang, while junction fragments featured one AGT overhang and multiple ACT overhangs. All structural hairpins featured AGT overhangs, whereas the amplification loop was designed exclusively with an ACT overhang. This is to drive each ligated DNA nanostructure to incorporates only a single amplification loop, preventing multiple amplification sites within a single assembly. The complementary strands comprising each fragment type were mixed separately and annealed to fold into respective DNA nanostructure fragments **(Figure 1B)**. The annealed fragments were then mixed and T4 ligation enzyme was added that joined the fragments via sticky-end ligation of the designed overhangs generating a complex library of interconnected structural assemblies. This process created a heterogeneous pool of nanostructures formed by the "gluing" of the individual components, forming a long single stranded ‘structure genome’ **(Figure 1C)**. To refine the library and remove unbound or partially ligated fragments, the ligated pool was subjected to Exonuclease III (Exo III) digestion. Exo III is a 3′→5′ exonuclease that specifically digests double-stranded DNA with free or recessed 3′ ends, while sparing DNA structures that lack accessible termini, such as circular fully ligated constructs. This selective degradation step effectively removed linear or incompletely ligated intermediates, which present free 3′ termini susceptible to Exo III attack. In contrast, fully ligated and topologically closed structures remained resistant to digestion, thereby enriching the pool for covalently closed and structurally complete nanostructures. The heterogeneity of the library and the impact of Exo III digestion were subsequently assessed by urea–PAGE and agarose gel electrophoresis. Following enzymatic treatment, several bands disappeared from the gel profiles, consistent with the selective removal of susceptible, non-protected assemblies **(Figure S1A)**. Finally, the structures from the ligated library containing the incorporated amplification loop were PCR amplified using primers specific to the loop, where one primer was produced with a 5’ phosphate modification for downstream enzymatic digestion. A gradient PCR across 50 °C–60 °C was used to find conditions that maintain library heterogeneity while minimizing non-specific amplification. A no template control (NTC) showed no product at any temperature, confirming that all observed bands were template-dependent. The successful PCR amplification of these structurally diverse templates generates a library of double-stranded DNA (dsDNA) structural genome. The ExoIII-treated library (Lib+ExoIII) amplified at all the tested temperatures however lower temperatures (50–55 °C) yielded distinct low molecular weight bands, whereas higher temperatures (58–60 °C) produced complex, smeared high-molecular-weight profiles, reflecting broader structural diversity **(Figure S1B)**. Based on these observations, 60 °C was selected as the optimal annealing temperature, as it yielded the most heterogeneous amplification profile. We next compared amplification between the undigested control library (Ctrl) and the Lib+ExoIII at 55 °C, 58 °C, and 60 °C. The Ctrl showed amplification at all temperatures but yielded fewer high–molecular-weight products whereas Lib+ExoIII produced more high–molecular-weight bands, indicating more heterogeneity in Lib+ExoIII amplified sample. In both cases, amplification efficiency and apparent product complexity increased with annealing temperature, with the most diverse profiles at 60 °C **(Figure S1C)**.

**Figure 1:**
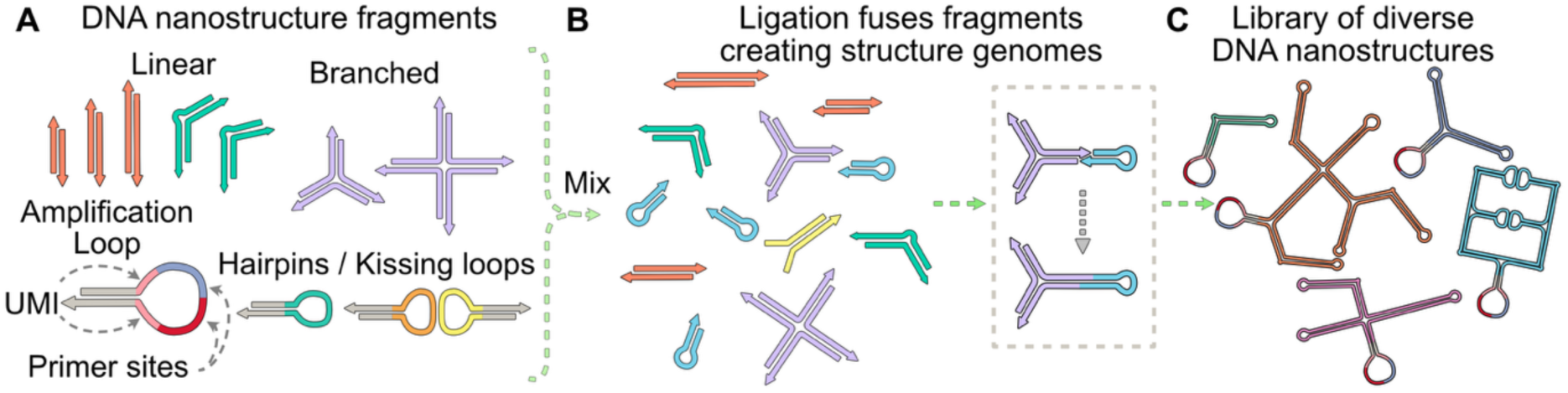
Schematic of generation and assembly of diverse DNA nanostructure library. (A) Small DNA nanostructure fragments of varying sizes and complexities linear, bulged, branched junctions, hairpins and amplification loop featuring primer-binding sites and twin 10-nt UMIs for lineage tracing. (B) The fragments are mixed and ligated via sticky ends to produce a diverse pool of DNA nanostructures (C) with many structures folding to form one single ‘genome strand’.

Following amplification, the library was purified to remove unincorporated primers, smaller fragments, and other reaction remnants resulting in the refined pool of assembled nanostructures. Next this library was sequenced that provided insights into the structural composition and complexity of the initial library, effectively profiling the initial mix of nanostructures. For subsequent cell selection experiments, the purified initial library was subjected to lambda exonuclease digestion. This enzymatic treatment selectively degraded strands with 5’ phosphate included in one of the PCR primers, rendering the nanostructures single-stranded. The resulting single-stranded structures were then subjected to a refolding process in a ligation buffer to refold them into their native conformations and subsequently used for cell selection assays. This structure library was imaged by AFM (atomic force microscopy), revealing numerous DNA nanostructures including some circular assemblies that may represent structures containing kissing loops **(Figure S1D)**.

### 2.2 Selection through repeated cellular uptake experiments

To investigate the structure-activity relationship and determine whether specific DNA nanostructures exhibit preferential cellular internalization, we utilized the structure library for cell selection experiments. We choose two cell models that are known to take up DNA: HEK293T, a human embryonic kidney cell line often used for transfection, and RAW264.7, a mouse macrophage-like cell line. A total of 2 µg of the DNA nanostructure library was separately incubated with in HEK293T and RAW264.7 cells for 4 hours to facilitate uptake. Non-treated cells were considered as negative control and were used as a comparative control to evaluate potential genomic or sub-cellular contamination. By analyzing their amplification profiles alongside those of the treated samples (cells exposed to nanostructures), we were able to verify that the sequences detected in downstream PCR originated from genuinely internalized nanostructures rather than background cellular material or processing artifacts. Following incubation, the cells were washed to reduce surface-bound nanostructures, allowing for the analysis of primarily internalized DNA. The cells were then lysed using RIPA lysis buffer and the resulting lysate was subjected to bead-based DNA purification to recover the DNA structures. Both treated and non-treated (control) purified lysate was subjected to agarose gel electrophoresis that revealed high molecular DNA bands that likely corresponds either genomic or some organellar DNA. Next upon PCR amplification of these lysates using primers targeting the amplification loop; a faint smear together with discrete bands appeared only in the treated sample that had been exposed to the DNA nanostructure library, whereas the control lysate yielded no detectable amplification product **(Figure S2A).** In order to verify that amplification was specific to our library, we performed in-silico PCR using UCSC genome browser BLAT tool against the human reference genome (GRCh38). No significant hits were identified, indicating that our primers do not non-specifically bind to host genomic DNA.^25^ We next tested PCR amplification for 10, 15, 20, and 25 cycles to determine an optimal cycle number that maximized yield while minimizing non-specific amplification. Although band intensity increased with cycle number, 15 cycles produced sufficient amplification and was therefore used for all subsequent lysate amplification experiments **(Figure S2B)**.

The amplified nanostructures were first purified using AmPure XP beads and then treated with lambda exonuclease to generate single-stranded DNA, followed by refolding in ligation buffer to reconstitute structured conformations. The refolded nanostructures were then reintroduced into their respective cell lines for the next round of selection. This iterative workflow of selection comprises of lysate purification, PCR amplification, purification, lambda exonuclease digestion, refolding, and re-incubation with the target cells was repeated for a total of ten cycles **(Figure 2A)**. A representative agarose gel illustrating the sequential processing steps for the round-6 lysate prior to round-7 selection as shown in **(Figure S2C)**.

**Figure 2:**
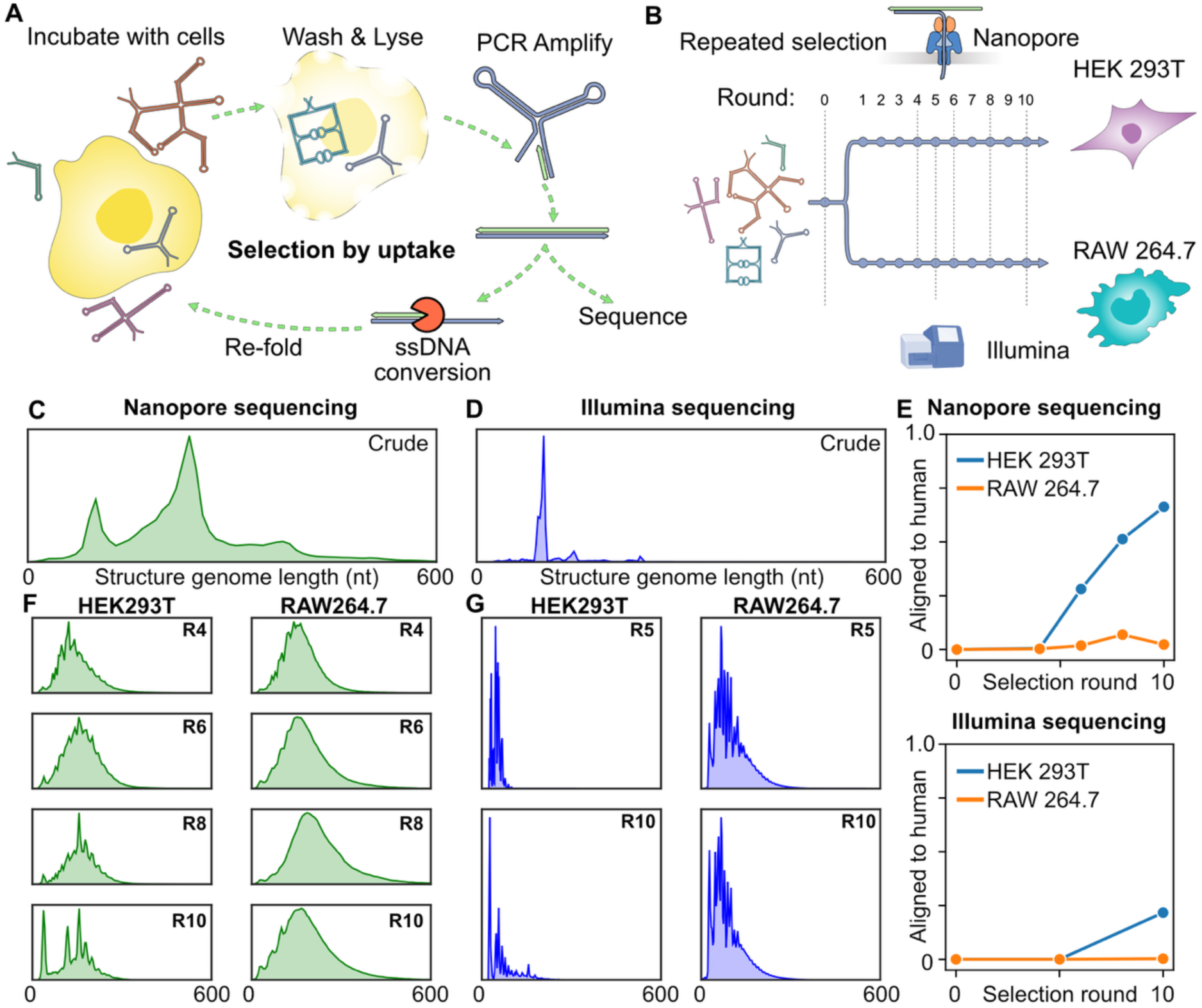
Selection of DNA nanostructures through cellular uptake. (A) Schematic of uptake mediated selection workflow for DNA nanostructures. DNA nanostructure library incubated with cells followed by washing and lysis to recover internalized nanostructures. These structures are then PCR amplified followed by ssDNA conversion using ƛ exonuclease and refolded for next round of selection. (B) This process is repeated and combined with high throughput sequencing to study the structural preferences of cellular uptake. The initial DNA nanostructure library was sequenced on (C) Oxford nanopore and (D) Illumina sequencing platforms. (E) Fraction of reads aligning to the human genome, indicating contamination from cellular DNA. (F-G) Structure genome length distribution over selection rounds in two cell lines using two sequencing methods after filtering out human genomic DNA reads.

To ensure that enrichment patterns is indeed due to cellular internalization rather than artifacts arising from PCR bias or preferential retention during purification, we performed an empty selection experiment in which the processed initial library was carried through 10 rounds of amplification and purification in the absence of any cellular selection step. Through this iterative enrichment process, we aimed to identify nanostructure candidates that consistently exhibited preferential internalization by HEK293T or RAW264.7 cells rather than any PCR amplification bias, thereby revealing insights into the structural features that drive cellular uptake.

In each round, aliquots of the amplified and purified dsDNA structure-genomes were saved for later high throughput sequencing to monitor the selection process **(Figure 2B)**. Oxford Nanopore sequencing (ONT) allows for rapid ligation sequencing with long read lengths at a relatively small scale, although with a lower read quality compared to other sequencing platforms, especially at the ends of the reads (where the UMI regions are located). The initial library as well as the selected libraries from rounds 4,6,8,10 in both cell lines were sequenced on flongle flow cells using a minion device, resulting in between 300k-900k reads per library **(Figure 2B)**.

To validate sequencing results from ONT the initial library as well as the selected libraries from round 5 and 10 in each cell line were re-sequenced using the Illumina sequencing platform with an error rate that is typically an order of magnitude lower, with the highest read quality at the ends and offers flow cells with significantly more reads than ONT. However, the longest commercially available Illumina read is 600 nt and the system has a strong preference for clustering and reading shorter fragments in pools with mixed read distribution.^26^ We prepared the libraries for Illumina sequencing by adding read primers and adapters via PCR. This resulted in between 6.2M - 8.7M reads per library **(Figure 2B)**. The read length distribution of the corresponding libraries is markedly different in the two sequencing methods. In the initial library, ONT sequencing shows a smooth distribution with peaks at 100, 240 and 370 nt, in contrast, Illumina sequencing shows an uneven distribution with peaks at 100, 140 and 240 nt where the peak at 100 nt dominates the sequencing **(Figure 2C and D)**.

A concern when amplifying DNA structures from cells is that we may also accidentally amplify genomic DNA from the cells. To detect potential genomic contamination these reads were aligned to the reference human genome GRCh38.^27^ As expected, we got no alignments in the initial libraries (R-0) as no human DNA should be present. This was confirmed by both nanopore and Illumina sequencing data for R-0 **(Figure 2E)**. The fraction of aligned sequences increases steadily in the HEK293T data sequenced by nanopore, up to around 62% of the reads in round 10 (R-10). The fraction aligned in the corresponding Illumina sequencing is lower, reaching 22 % in R-10 of selection in HEK293T, this could be due to the size preference of Illumina sequencing disfavoring longer fragments of genetic origin. In contrast, the fraction of reads mapping to the human genome in the RAW264.7 cell line remained below 7 % in all rounds and both sequencing methods. This may be due to the more promiscuous macrophage cells taking up DNA more effectively leading to more target DNA in the cell lysate and less unspecific amplification. Still, even though the sequencing round mostly contaminated with human DNA we still got 80k reads for analysis of DNA nanostructure uptake in cells.

After filtering out reads not from DNA nanostructures the preference for structure sizes in the two cell lines can be seen in the change of the distribution of genome lengths. In ONT sequencing of the cell uptake in HEK293T cells a transition can be seen from the smooth distribution in the initial library to a shorter average size with several pronounced read length spikes **(Figure 2F)**. A similar effect can be seen in the Illumina sequencing although with a stronger preference for shorter sequences **(Figure 2G)**. Interestingly the selection in RAW264.7 cells show a more modest change in length distribution in both sequencing methods compared to the initial distribution further indicating that macrophage cells promiscuously uptake DNA structures with little size preference.

### 2.3 Structural preferences in cellular uptake

The sequencing data derived from the initial library and evolved library pool after successive rounds of selection allows for the analysis of both general structure characteristics that increase cell uptake and the identification of individual nanostructures that are evolutionarily favored. Each DNA nanostructure genome is a combination of ligated fragments, terminated at either end by 10 nucleotide UMI regions and the primer binding sites from the amplification loop **(Figure 3A).** By decoding the genome fragment composition, we can understand what type of structure features lead to preferential uptake in certain cell types. Although nanopore sequencing has a higher error rate, it is possible to reconstruct the fragment composition of the structure genomes using the tool concept that we recently developed.^28^ By calculating the composition of structures and relative abundance of fragments in the initial library and comparing it to later sequencing rounds we can calculate the change in abundance of each fragment type and use it to infer evolutionary fitness benefits of structures including certain fragments. This revealed a clear preference for linear fragments in both cell lines as well as a strong negative selection for kissing loop segments **(Figure 3B)**. The relative changes were larger in the HEK293T selection experiments than in the RAW264.7 again indicating that the macrophage cell line is more promiscuous in uptake. The concert analysis also allowed us to look at the ordering of fragments in the structure genomes, surprisingly many structures did not contain the theoretically expected ordering of fragments indicating that ligation does not occur as expected. Off-target ligation of the non-palindromic toehold could lead to the ligation of non-hybridized fragment oligonucleotides or the incorporation of multiple amplification loop, creating a higher-than-expected diversity of structures in the library.

**Figure 3:**
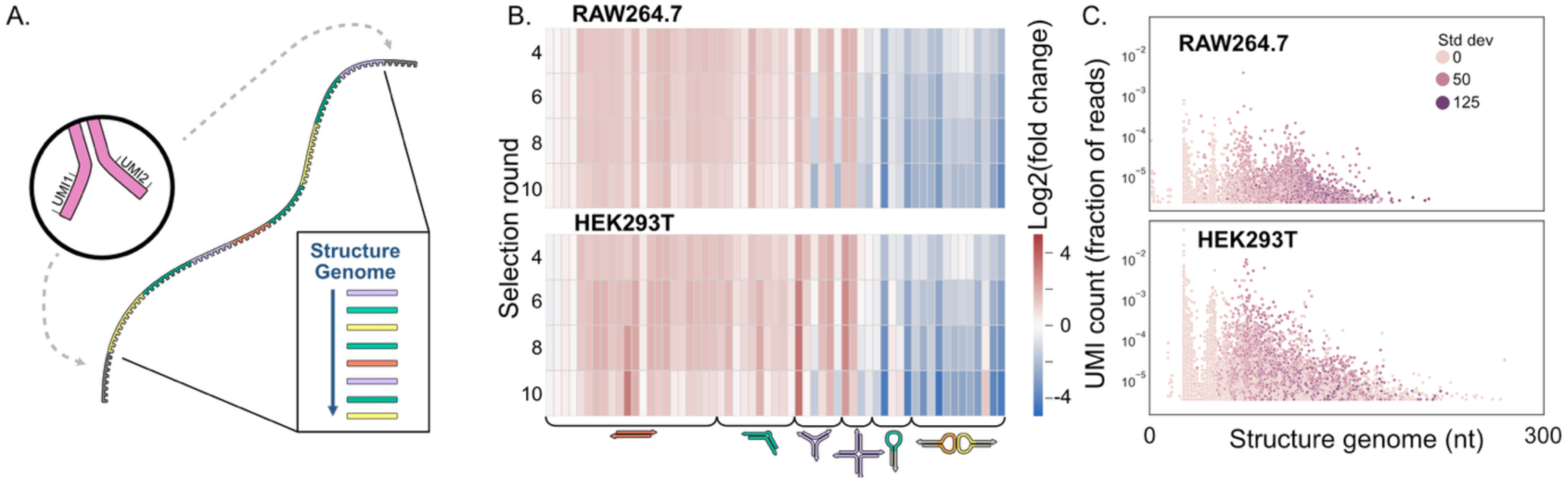
Analysis of sequenced DNA nanostructure genomes. (A) Each structure genome is comprised of ligated constituent fragments and terminated in either end with a 10 nt unique molecular identified (UMI). (B) Change in relative abundance of structure fragments compared to the initial library in two cell lines, from ONT data. (C) Distribution of abundance, genome size (excluding primer sites) and length variability of UMI labeled structure genomes after 10 rounds of selection in two cell lines, from Illumina data.

Each structure genome is labeled with one 10 nit UMI at either end of the genome, together these two UMIs result in a 20 nit barcode for each unique structure formed at the initial library generation. The underlying diversity of 4^20^ ≈ 10^12^ means that it’s unlikely for several structures to get identical barcodes. The analysis of structure UMI’s requires the comparison and identification of identical 20 nt segments between reads where read errors are likely to lead to miss-identification. ONT sequencing has an error rate that can be above 10% in the start of reads making the accurate determination of UMI identity somewhat difficult. In comparison Illumina sequencing typically offers an error rate below 0.1% in the start of reads, making it suitable for the identification of re-occurring UMIs. Plotting the abundance of individual UMI’s together with the corresponding genome length after 10 rounds of selection reveals thousands of individual DNA nanostructures that have increased in number. This effect is clearer in HEK293T cells with 238 UMIs appearing more than 1000 times in sequencing compared to just 11 UMIs in the RAW264.7 selection experiments indicating a more stringent selection process in the HEK293T cells **(Figure 3C)**. The difference in genome length in structures with identical UMIs is illustrated by plotting the standard deviation of the extracted genomes, for many UMI’s the standard deviation is 0 but for some it’s significant indicating that structure genomes have experienced divergent mutations during the selection process.

In our control experiment where we performed 10 rounds of ‘empty’ selection, the sample from round 5 (R-5) and round 10 (R-10) were sequenced using ONT platform. This data revealed no strong preference for any UMI, in the sequencing data after 10 rounds the top UMI occurred only in 0.02% of reads and around 97% of reads are labeled with unique UMI’s. The empty selection resulted in relatively smooth length distribution with higher average genome lengths than either of the cell selections **(Figure S3A).** Similarly to the cell selection, loops were negatively selected for although the changes were smaller in the empty selection **(Figure S3B)**.

### 2.4 Identification and production of DNA nanostructure candidates from sequencing data

#### 2.4.1 Selection from nanopore data

Nanopore sequencing showed enrichment of some candidate sequences that were progressively enriched across successive selection rounds. This prompted us to further investigate these candidates to understand the basis of their preferential accumulation. At this point, it remained uncertain whether this enrichment reflected genuine cellular selection pressures or resulted from stochastic events or methodological biases inherent to the process. In our initial analysis, we prioritized sequences longer than 200 nucleotides and further filtered for candidates with high UMI counts after selection. Based on their strong enrichment profile and consistent growth across selection rounds we initially selected four candidate structure genomes of lengths 208 bp-H5A, 283 bp-H5B, 375 bp-H5C, and 485 bp-H5D from HEK293T round-5 nanopore data. These candidates exhibited distinct enrichment dynamics over the 10-round selection cycle. H5A and H5C showed sustained exponential enrichment through Round 10, with H5A displaying the most robust increase. In contrast, H5B and H5D stagnated following Round 8 **(Figure S4A)**.

To validate this enrichment profiles and produce these structures individually for further testing we designed structure-specific primers targeting UMI regions at the start and end of the structure genomes of interest, with the reverse primers labeled with a Cy5 fluorophore. These primers allowed us to selectively amplify each candidate structure from purified lysates collected across different selection rounds to further validate their enrichment profile. Using PCR-amplified and purified Round-5 lysate as the template, we tested different annealing temperatures ranging from 45°C to 50°C based on the annealing temperature of primer pairs and successfully amplified all four candidates, confirming their enrichment. We imaged the gel in both SYBR Safe **(Figure S4B)** and Cy5 **(Figure S4C)** channels to verify specific amplification and incorporation of the Cy5-labeled primer. We observed a single discrete band for each candidate, without any non-specific amplification in both channels and minor signal from unincorporated primers at the gel bottom in Cy5 channel. We next purified these PCR-amplified candidates using AmPure beads and the SYBR Safe channel revealed discrete bands for the purified structures **(Figure S4D)**, while the Cy5 channel **(Figure S4E)** confirmed the removal of free, unincorporated primers from the gel bottom, ensuring high purity for further processing.

These structures were selected before we were aware of and filtered out structure genomes that aligned with human genomic DNA. The subsequent analysis revealed that these highly enriched sequences comprised a mixture of genomic fragments and synthetic elements originating from the initial library. Although the precise mechanism underlying their cellular processing remains unclear, these sequences exhibited exponential accumulation across successive selection rounds.

We next determined the genomic content of these highly enriched candidates, we mapped these candidates to the human reference genome (GRCh38) using the Blast-like Alignment Tool (BLAT). Alignment analysis revealed that all analyzed candidates produced multiple alignment hits with variable score distributions. The top five hits for each candidate, along with their detailed alignment maps, are presented in **(Figure S5A**). Notably, across all analyzed candidates, alignments were localized to the 5’ or internal segments of the query, with no significant alignment observed at the 3’ terminus **(Figure S5B**).

To further investigate the extent of genomic contamination, we performed PCR amplification of each candidate from the initial (pre-selection) library R-0, empty selection library R-5 **(Figure S6A)** and R-10 **(Figure S6B)**, and non-treated purified cell lysate **(Figure S6C)**. We observed that the initial library R0 showed all candidates with a faint H5D signal; R5 showed strong signals except for H5D, while R10 showed reduced H5C and no H5D. The HEK293T control showed signal for H5A and a faint H5B. We performed an initial series of cell-uptake assays to characterize the intracellular behavior of each individual structure. Although later analysis showed that the H5A and H5B candidates contain genomic fragments, their pronounced enrichment warranted further study. Using templates from selection round 5, we monitored the uptake of H5A, H5B, H5C and H5D Cy5 fluorescence was substantially higher in treated cells than in untreated controls, with the H5B and H5C showing more diffusive and homogenous fluorescence compared to H5A and H5D**(Figure S7A)**. By round 10 there was significant shift with H5B showing the brightest signal compared among all the samples. The initial library showed no visible fluorescence signal compared to R-10 library which has some fluorescence signal but weaker than the H5B highlighting the significance of enrichment of library and selected candidates in the cellular uptake **(Figure S7B)**. To exclude the possibility that the fluorescence signal originated from the Cy5-labeled primer alone rather than from amplified products, we performed a primer-only control in which the forward primer and Cy5-labeled backward primer were mixed and incubated with the cells. These samples showed minimal fluorescence, further confirming that the observed signal arises from the corresponding amplified structures **(Figure S8)**. Subsequently the candidates were evaluated for internalization in both HEK293T and RAW cells to assess potential preferences for sequences selected in a specific cellular context.

#### 2.4.2 Selection from Illumina data

We next selected candidate sequences from both RAW264.7 and HEK293T datasets generated by Illumina sequencing, given the higher read accuracy of Illumina data. Candidates were chosen primarily from round 10 samples of both cell lines. At this stage, long double-stranded DNA fragments (up to 600 bp) corresponding to these candidates were synthesized as single contiguous constructs by ANSA Biotechnology. To enable downstream amplification and fluorescence-based tracking, primer-binding sites were incorporated at both the 5′ and 3′ termini of each construct. This design enabled selective PCR amplification using Cy5-labeled primers for subsequent cellular uptake assays. Candidate selection was designed to capture enrichment patterns from both cell lines and multiple selection rounds. Since earlier candidates had been derived predominantly from HEK293T selections, we expanded our analysis to include RAW264.7–derived sequences. Four candidates were selected from RAW264.7 round 10 (R10A, R10B, R10C, and R10D), based on their high UMI counts and absence of genomic DNA. In parallel, four candidates from HEK293T round 10 Illumina data were selected and labeled H10A, H10B, H10C, and H10D using the same selection criteria. In addition, we applied two strategies to select control candidates for experimental comparison. First, we selected four candidates that appeared in the Illumina sequencing data of the initial library but were completely absent from Illumina sequencing data of the selected libraries, this we called Crude Control A-D. Secondly we applied random selection from Illumina data of the initial library to find 9 candidates of different length, called Random 1-9. Sequence identities and corresponding UMI counts for all candidates are provided in **(SI Table 4)**.

The integrity and purity of the synthesized fragments were first verified by gel electrophoresis, which confirmed the presence of a single discrete population for each construct **(Figure S9A)**. These fragments were subsequently used as templates for PCR amplification. The amplification of ANSA-synthesized candidates selected from HEK293T and RAW264.7 selections proceeded robustly and yielded single discrete bands after amplification **(Figure S9 B&C)**. However, the initial amplification of random and crude control sequences proved more challenging. Although these control templates appeared as single populations prior to amplification, PCR reactions frequently produced multiple bands indicating nonspecific amplification for these constructs **(Figure S10A, B&C)**. We tried optimizing the PCR conditions and observed that elevated primer and template concentrations might have contributed to nonspecific amplification. Accordingly, the initial denaturation time was extended from 3 to 5 minutes, primer concentration was reduced significantly from 0.4 µM to 0.1 µM, and template concentration was lowered from 0.1 nM to 0.01 nM. These adjustments effectively eliminated nonspecific products and enabled clean amplification of all candidate and control sequences **(Figure S10D)**. Following optimization, a total of fourteen candidates were selected for subsequent cellular uptake studies. These included enriched sequences derived from both HEK293T and RAW264.7 selections, as well as non-enriched random sequences serving as negative controls. From the HEK293T selection, candidates H10A, H10B, H10C, and H10D were chosen, while from the RAW cell round 10 selection, candidates R10A, R10B, R10C, and R10D were selected. In addition, non-enriched random sequences (Random1, Random4, Random6) and the Crude-control-C were included as negative controls. The H5B and H5D candidates from the initial screen were also synthesized independently (ANSA) and included in the study. We also verified the selected candidates before cell stimulation via gel electrophoresis and observed a single predominant population as shown in **(Figure S11).** The Secondary structure prediction of the finalized candidates performed using mfold^29^ depicts a diverse structural spectrum where the most stable conformations of selected candidates displayed higher prevalence of loops whereas the randomly selected candidates displayed distinctly longer double-stranded stems **(Figure S12)**. The thermodynamic stability and other structural characteristics of these candidates were evaluated based on mfold prediction data with results summarized in (**Table S5).**

### 2.5 Cellular profiling of candidates by flow cytometry and microscopy

Following the final selection of candidate structures from both cell lines, we assessed and compared their cellular internalization as purified individual nanostructures. We have a total of fourteen candidates, and each candidate was amplified with fluorophore labelled primers, purified, subjected to lambda exonuclease digestion, and refolded according to the protocol described above. The refolded structures were incubated with cells at a final concentration of 25 nM for 1 hour and analyzed using flow cytometry **(Figure 4A)**. Given that these candidates were evaluated as purified individual structures, internalization was expected to occur more efficiently than in pooled library conditions.

**Figure 4:**
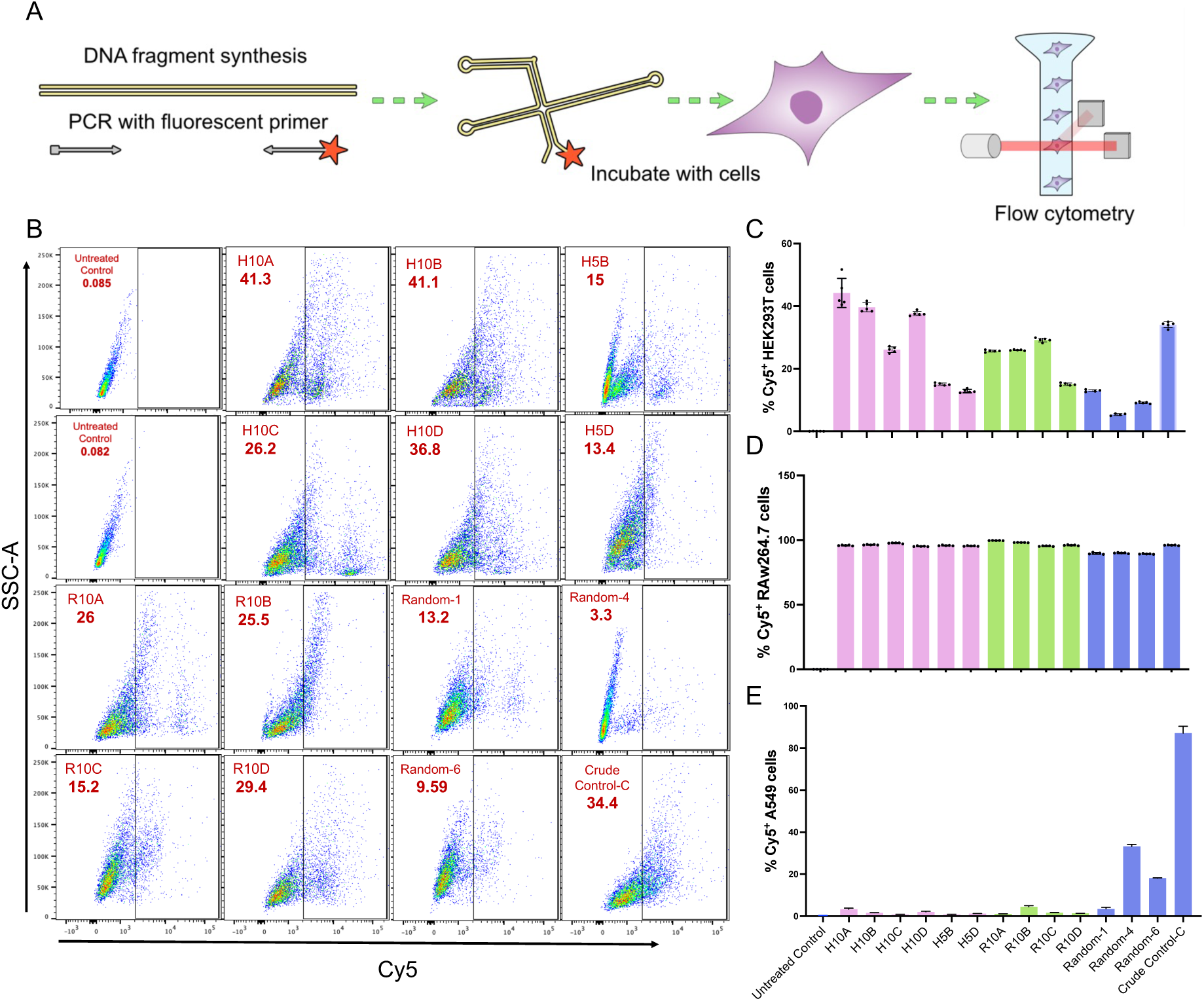
Quantification of DNA nanostructure internalization via flow cytometry. (A) Schematic illustrating the flow cytometry workflow, in which each candidate nanostructure was amplified using Cy5-labeled primers, incubated with cells, and subsequently analyzed using flow cytometer to quantify cellular uptake. (B) Representative flow cytometry scatter plots of selected candidates showing Cy5+ cells in HEK293T cells. (C) Quantification of Cy5+ HEK293T cells as a percentage of total cells in each sample group. (D) Quantification of Cy5+ RAW264.7 cells as a percentage of total cells in each sample group. (E) Quantification of Cy5+ A549 cells as a percentage of total cells in each sample group.

We performed flow cytometry with the finalized candidates to quantitatively evaluate DNA uptake at single-cell resolution. This method provides a broader quantitative assessment by recording multiple events per sample allowing robust statistical comparison of uptake efficiencies across different treatment conditions. For each sample, 10,000 events were acquired, and all measurements were performed in triplicate or quadruplicate. Cy5 fluorescence was detected in the APC channel.

In HEK293T cells, the untreated control population remained entirely confined to the left region of the APC histogram, indicating negligible background signal. In contrast, all treated samples exhibited a rightward shift, consistent with cellular internalization of DNA nanostructures. **(Figure 4B)**. Among the tested candidates, H10A and H10B showed the largest Cy5-positive cell populations, followed by H10D and the Crude-control-C **(Figure 4C)**. Notably, candidates derived from the enriched cell line demonstrated higher levels of internalization compared with those selected from the RAW cell line. Interestingly, Crude-control-C exhibited a substantially larger Cy5-positive cell population than most randomly selected candidates, highlighting an inherent limitation of the selection strategy whereby potentially effective candidates may not be enriched or identified.

In the RAW264.7 macrophage like cells, treatment induced a pronounced rightward shift of the entire cell population, reflecting the intrinsically high and promiscuous uptake capacity of this cell line. Utilizing this gating strategy >90% of the population were Cy5-positive **(Figure 4D)**. To better resolve differences among candidates, a Cy_5_^high^ gate was applied to identify strongly positive cells. Within this gated population, R10B yielded the largest Cy5-positive cell population, followed by Crude-control-C and R10A, whereas randomly selected candidates exhibited the smallest proportion of Cy5-positive cells **(Figure S13)**.

To study the uptake behavior of structures in a cell line not used in selection we used the human lung cancer cell line A549. Compared to the other cell lines the overall DNA uptake was low. In flow histogram for selected candidates the majority of cell population is fixed towards the left and very few cells move towards the right whereas control candidates induced a greater rightward shift **(Figure S14)**. Crude-control-C exhibited the highest proportion, followed by Random-4 and Random-6. All remaining candidates resulted in fewer than 10% Cy5-positive cells, indicating limited internalization in this cell line **(Figure 4E)**.

We next employed Laser scanning confocal microscopy to determine whether the DNA nanostructures were merely surface bound or truly internalized. Optical sectioning was performed for each sample and only the central z-slices were used to generate representative images, thereby minimizing contributions from surface associated fluorescence. All samples were imaged under identical acquisition settings to permit comparison of fluorescence intensity across cell types and candidates.

In HEK293T cells, uptake was highly heterogeneous: a subset of cells exhibited intense fluorescence throughout the cytoplasm and nucleus, whereas others displayed sparse, punctate signals. Candidate H10A showed clear nuclear localization, while H10B, H10C, and H10D were confined predominantly to the cytosol; notably, H10C and H10D produced markedly weaker fluorescence than H10A and H10B. The H5B and H5D showed more homogenous and diffusive cytosolic signal **(Figure 5A)**. Some of the RAW cell selected candidates like R10B and R10C seemed to show nuclear signal. The signal from R10B was only nuclear however it looked like the bleed through of Hoechst signal. But all the images were acquired under sequential and identical setting so this nuclear signal could indeed be due to the candidate. The R10C and R10D showed heterogenous signal like H10 candidates **(Figure 5B)**. Surprisingly the randomly selected control candidates showed hardly any signal **(Figure 5C)**.

**Figure 5:**
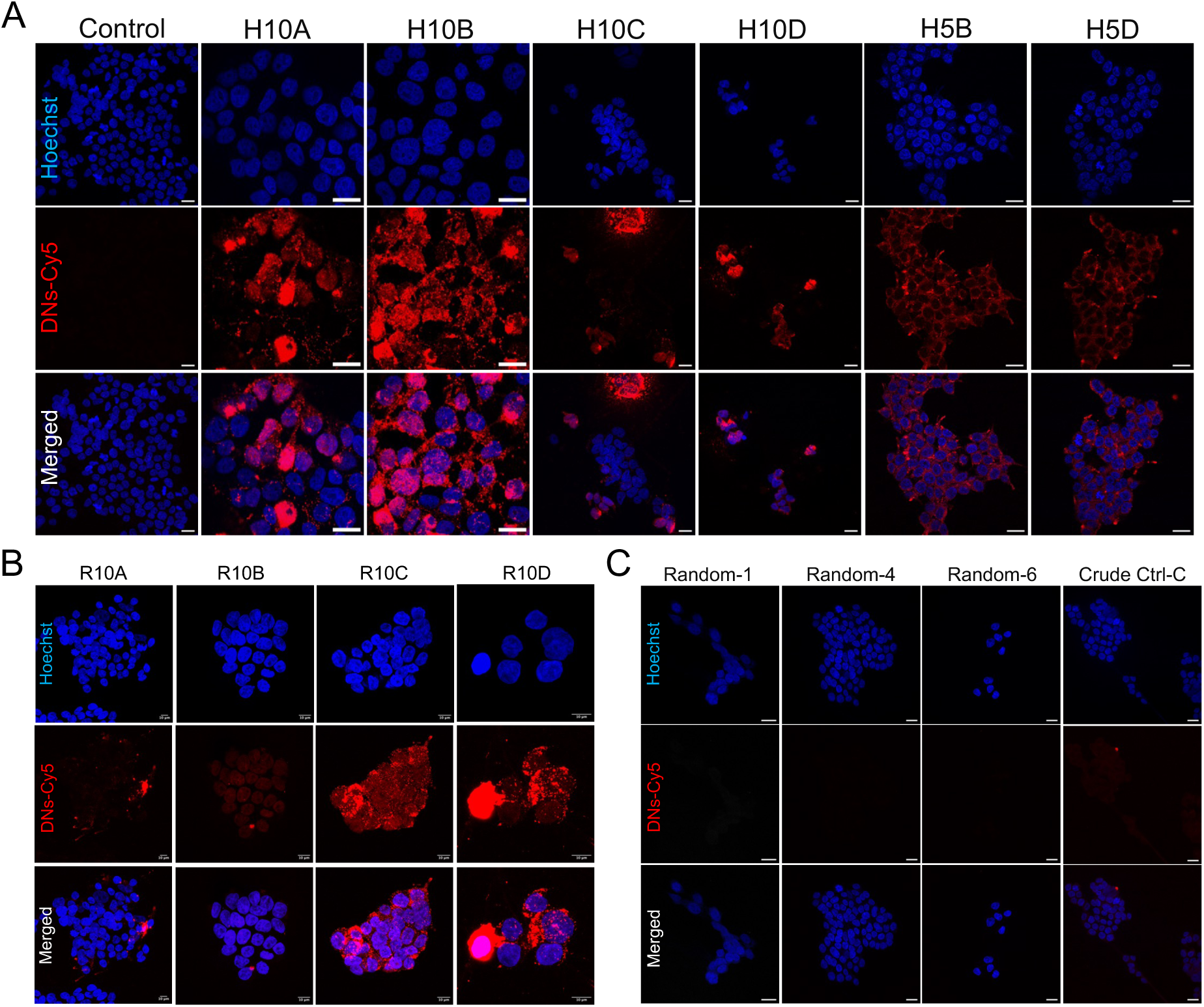
Qualitative assessment of cellular uptake of final candidates in HEK293T using laser scanning confocal microscopy. (A) Confocal images of cells treated with HEK293T-selected DNA nanostructure (DN-Cy5). (B) Confocal images of cells treated with RAW cell-selected DNA nanostructure. (C) Confocal images of cells treated with randomly selected DNA nanostructure. For all panels, the red channel represents DN-Cy5 uptake, and the bottom panels represent merged images of all channels with nuclei stained with Hoechst dye (blue). Scale bar: 20 μm for A and C and 10 μm for panel B.

In comparison, RAW 264.7 cells demonstrated more uniform internalization; cells within a single focal plane presented comparable signal intensities. All DNA nanostructures remained cytosolic with no detectable nuclear signal. Certain candidates formed distinct punctate structures suggestive of endosomal sequestration, whereas others displayed a diffuse cytosolic pattern **(Figure S15)**.

In A549 cells, overall fluorescence intensity was visibly reduced compared to other cell lines under identical acquisition parameters. The signal intensity was visibly lower in H5B, H5D, and H10D among candidates selected from HEK293T. Furthermore, candidates selected from RAW264.7 exhibited weaker uptake compared to the HEK-selected group. Interestingly, while the majority of random controls showed negligible internalization, the Crude-control-C displayed significantly higher fluorescence intensity than any other candidate in A549 cells. This was similar to what we observe in the flow cytometry data where Crude-control-C showed the highest number of Cy5 positive cells compared to rest of the candidates **(Figure S16).**

These observations reveal cell-type dependent differences in internalization efficiency and subcellular distribution of the DNA nanostructure candidates. The pronounced heterogeneity and occasional nuclear entry in HEK293T cells contrast with the more homogeneous, cytosol restricted uptake in RAW 264.7 cells. The low fluorescence in A549 cells may reflect reduced uptake capacity of the constructs in the non-selected cell line. The elevated signal from the Crude-control-C underscores the limitation of the method that some non-selected candidates that were lost from the selection pool might have some interesting behavior.

## 3. Discussion

Developing structures that can specifically target and deliver drugs to certain cell types is a major goal of DNA nanotechnology. The strength of DNA nanotechnology is the ability to precisely control size, shape and mechanical properties to optimally achieve a target function. Although several recent studies have demonstrated that DNA nanostructure design choices greatly influence uptake in different cell types^7,9,30,31^, the optimal combination is unknown and current methods limited to testing structures one by one.

Here we attempt to invert the process by using cell uptake as a selection pressure to discover structures that are preferentially taken up by cells from a large initial library. In order for selection to function, the DNA structures must be compatible with amplification and sequencing, this is not possible with DNA origami and tile techniques where the composite strands disassemble when heated. Instead, we base our approach on DNA nanostructures that fold from one long single strand that we achieve by randomly ligating smaller DNA nanostructure fragments assembled from 1-5 oligonucleotides. Although this limits the number of possible structural motifs, DNA nanostructures that fold from a single long strand are easier to scale up than other approaches as the single genome can be amplified enzymatically.

The underlying assumption that cells preferentially uptake DNA nanostructures with certain shapes and mechanics appears to vary highly among cell types. In this first proof-of-concept work, we chose cell lines that are known to abundantly take up DNA: a macrophage-like cell line (RAW) and a cell line well known for its ease of transfection (HEK293T). Using these cells, we demonstrated that DNA nanostructures could be amplified, sequenced and refolded for repeat selection from cell lysate. However, the macrophage-like cell appeared very promiscuous in their uptake preferences making them an important control but not ideal for discovering cell specific targeting. This was true to a lesser amount in the HEK293T cell where we were able to discover and select for DNA nanostructures that were taken up more effectively than randomly selected DNA nanostructures. Intriguingly, we tested our monoclonal DNA nanostructure candidates for uptake in a cell line not used in the selection: the human lung cancer cell line A549 and it showed a great variety of uptake preference of fluorescently labeled structures indicating that cancer cell lines may be a good target for structure type selection.

In any synthetic selection technique, it is important to be mindful that results may be a combination of several parallel selection pressures. Although we primarily discuss cell type specific uptake here, the selection process we see is likely to also be influenced by the stability of the DNA structures in the cell culture: structures that degrade are less likely to be amplified and sequenced something that is a positive property of a DNA nanostructure for drug delivery. This could be explored further by parallel selection experiments where the incubation time is varied. PCR bias, where some structures are more preferentially amplified, is a more concerning selection pressure as it could create the illusion of cell-selection. Here we have performed control experiments where we only amplify the structures without cellular incubation and not seen any clear trends, but this is still an important factor to keep in mind. In our results we see a structure (Crude-control-C) that is not enhanced for selection in either cell line but still appears to be effectively taken up, this may be due to the abovementioned factors (poor stability or amplification) or may just be a random effect when the selection pressure is not strong.

Our sequencing results after 10 rounds still contained a diverse set of DNA nanostructures that have been enhanced by a mixture of selection pressures. It is only feasible to individually test a handful of these structures and knowing what candidates to pick is not trivial. Future computational work may be able to deconvolute the parallel selection pressures for example by comparing multiple selection experiments or using structure prediction at scale to group structures with similar features. This may allow us to find better candidates, either in new experiments or in the sequencing data we publish here.

In this work we only explore the effect of the size, shape and mechanics of DNA nanostructures on cellular uptake preferences. It is known that the addition of targeting groups greatly influences the cell type specific targeting of structures.^4,32^ In future work, DNA aptamers could be directly integrated in DNA loops and used to target surface groups on target cells increasing the selection pressure. Aptamers integrated in the structure genome could be directly amplified for repeat selection and sequencing. Other functional groups such as proteins or peptides cannot be directly integrated in the genome but could be conjugated to DNA handles complementary with loops in the DNA structures and be added to the library in each selection round.

The structures selected for in this study could either stick to the cell surface or be taken up inside the cells by several mechanisms. It is feasible to use the tools developed here to study the uptake mechanisms in more detail, for example by chemically blocking certain uptake routes^33–35^ or use cell fractionation in each round to select for structures that end up in certain cellular compartments. It is conceivable that this type of selection could even be used in vivo where a library is injected into an animal followed by selection from certain tissue types or tumors.

In conclusion we have developed a method for turning DNA nanostructure cell targeting from a design to a selection process. We have developed a new type of DNA nanostructure compatible with amplification and sequencing and developed the processes needed to use cellular uptake as a selection pressure. We have demonstrated that there is cellular preference for certain types of DNA nanostructures, but the specificity is greatly dependent on the cell type. We hope that this can be a tool to discover DNA nanostructures that would be intractable to find with current methods to potentially target diseases lacking efficient therapies today.

## 4. Material and Methods

### 4.1 DNA library preparation

We employed a library of orthogonal 20-nucleotide sequences as the foundation for designing DNA fragments with diverse geometries and sizes. These sequences were optimized to have similar melting temperatures, low cross-interaction and no homopolymer repeats. From these we created fragments including linear duplexes of varying lengths (10–20 nt), bulged fragments with bulge sizes ranging from 1–5 nt, hairpin structures (10–18 nt), kissing loops (10–20 nt), three-way junctions, four-way junctions, four-way junctions containing a thymine (T) at the junction. We also create an amplification loop hairpin with two adjacent primers binding sites flanked by 10 nt random sequence region that function as Unique Molecular Identifiers (UMI). The design and structural modelling of these fragments were performed using oxView^36^ oxDNA^37^, a computational tool for DNA nanostructure simulation. DNA oligonucleotides were purchased from IDT at a stock concentration of 100 µM and diluted to 10 µM in nuclease free water for further use. Complementary strands for each DNA fragment were annealed in 1×T4 DNA ligase buffer using a temperature ramp from 90 °C to 4 °C over 30 min. For each fragment, 45 µL of the 10 µM oligonucleotide mixture was combined with 5 µL of 10× DNA ligase buffer, yielding an effective concentration of 9 µM per fragment. All 37 annealed fragments were then pooled together in equimolar amounts and mixed with T4 DNA ligase to generate the ligated library. This was incubated for 10h at 16°C, followed by an overnight hold at 4°C

#### 4.1.1 Exonuclease Treatment and PCR amplification

Following ligation, the library was treated with Exonuclease III (Exo III; NEB, 100 U/µL), which selectively digests double-stranded DNA from 3′ to 5′ at linear or nicked sites. Because Exo III progressively resects dsDNA—producing intermediates with short single-stranded extensions and ultimately ssDNA—this step preferentially removes unligated or partially folded fragments while preserving stable, fully assembled structures. Exonuclease digestion was performed using the following temperature program: 37 °C for 30 min, followed by 70 °C for 20 min to heat-inactivate the enzyme. A negative control lacking enzyme was processed in parallel. Reactions were set up as follows:

| Sample Name | Sample Vol. (μl) | Exo-III (μl) | MilliQ (μl) | Total Vol (μl) |
| --- | --- | --- | --- | --- |
| Ligated Mix | 95 | 5 | 0 | 100 |
| Negative Control | 45 | 0 | 5 | 50 |

The Exonuclease III–digested library served as the template for PCR amplification using primers specific to the UMI-containing amplification loop. In principle, only structures that had successfully incorporated the primer loop should be amplified under these conditions. To optimize amplification while minimizing nonspecific products, we tested annealing temperatures ranging from 55 °C to 65 °C and used primers at a final concentration of 0.4 µM. A template concentration of 1 ng/µL was used, and multiple cycle numbers (10, 15, 20, and 25 cycles) were evaluated to avoid overamplification, nonspecific products, or primer depletion. The PCR reaction setup for a 50 µL reaction volume is shown below.

| Component | Volume (μl) |
| --- | --- |
| 2x DreamTaq Master Mix | 25 |
| Fwd primer (10 μM) | 2 |
| Bwd primer (10 μM) | 2 |
| Lib+ExoIII | 2 |
| H<sub>2</sub>O | 19 |
| Total Vol (ul) | 50 |

Based on these optimization experiments, 60 °C was selected as the annealing temperature and 15 cycles as the optimal amplification regime. PCR reactions were prepared using DreamTaq 2× Master Mix (Thermo Fisher Scientific) in 20–25 µL reaction volumes. The thermal cycling protocol consisted of an initial denaturation at 95 °C for 2 min, followed by 15 cycles of 95 °C for 30 s, 60 °C for 30 s, and 72 °C for 30 s, with a final extension at 72 °C for 10 min. Following amplification, products were purified using AmPure beads to remove residual primers and small contaminants.

#### 4.1.2 AmPure Purification Protocol

PCR amplicons were purified using solid-phase reversible immobilization (SPRI) with AMPure XP paramagnetic beads (Beckman Coulter). Beads were added to each sample at a 1.8× bead-to-sample volume ratio (324 µL of thoroughly resuspended beads to 180 µL of PCR product) and mixed on a rotary mixer for 5 min at room temperature to allow DNA binding. Samples were placed on a magnetic stand, and the supernatant was discarded once bead separation was complete. The bead-bound DNA was washed twice with 200 µL of freshly prepared 75% ethanol. Beads were briefly air-dried to remove residual ethanol while avoiding over-drying. Purified DNA was then eluted in 100 µL of nuclease-free water and subsequently processed either for nanopore sequencing or for λ exonuclease digestion and refolding prior to cell selection assays.

#### 4.1.3 Lambda exonuclease digestion and refolding

To prepare the library for subsequent rounds of cell selection assays, the purified dsDNA library was subjected to lambda exonuclease digestion to convert the library into ssDNA. Lambda exonuclease (NEB; 5 U/µL) is a 5′→3′ exonuclease that specifically binds to 5′-phosphorylated double-stranded DNA (dsDNA). Upon binding, the enzyme selectively digests only the phosphorylated strand, progressing unidirectionally from the 5′ end. As nucleotides are removed, the complementary strand remains intact, generating long single-stranded DNA (ssDNA) that facilitates downstream refolding behavior. Following the manufacturer’s recommendation-1 unit of enzyme per ∼1 µg of purified dsDNA, the digestion reactions were assembled based on Qubit-quantified dsDNA concentrations. The reaction was incubated at 37°C for 30 minutes, followed by heat inactivation at 75°C for 10 minutes to terminate enzymatic activity. Post-digestion Qubit measurements confirmed that most of the library had been converted to ssDNA: the dsDNA signal was substantially reduced, whereas the ssDNA concentration increased, opposite to the pre-digestion profile. This shift is consistent with efficient strand-specific degradation by lambda exonuclease.

The digested library was then refolded in DNA ligase buffer using a controlled temperature ramp to promote recovery of the nanostructures intended secondary conformations.

### 4.2 AFM sample preparation and imaging

Samples were diluted 5-fold in AFM buffer (1X TE buffer supplemented with 10mM MgCl_2_). 10 μL of diluted sample solution was applied to freshly cleaved mica and allowed to incubate for 30s. 4μL of 5mM NiSO_4_ was added to the mica on top of the sample and allowed to incubate for another 4min 30s. The mica was then washed by applying and removing 1 mL of AFM buffer using a pipette. After washing, 1.5L of AFM buffer added to the mica for imaging. AFM imaging was performed in liquid using a JPK Instruments Nanowizard 3 Ultra with a Si_3_N_4_ Olympus Biolever Mini cantilever in a Bruker Hyperdrive glass block. Images were taken in AC Mode with a frequency in the range 25-35kHz and a setpoint of 70-80%.

### 4.3 Cellular Selection in HEK293T and RAW264.7

We conducted cellular selection experiments using two cell lines: HEK293T and RAW264.7. Both cell lines were cultured in DMEM supplemented with 10% fetal bovine serum (FBS) and 1% penicillin-streptomycin. Cells were maintained at 37°C in a humidified incubator with 5% CO₂. For subculturing, HEK293T cells were detached using trypsinization and reseeded following standard protocols. In contrast, RAW264.7 cells, being adherent macrophages, were harvested using a cell scraper and subsequently sub cultured into fresh medium.

#### 4.3.1 DNA library treatment and uptake protocol

Cells were seeded in six-well plates one day prior to the experiment to ensure adequate adherence and optimal confluency. The following day, 2 µg of the DNA library sample was added to each well in serum-free DMEM to facilitate cellular uptake without serum interference. Cells were incubated with the DNA library for 4 hours.

#### 4.3.2 Post-Incubation Processing

After incubation, cells were washed three times with 1x PBS to remove any surface-bound DNA structures. For HEK293T cells, washing was performed gently to avoid dislodging the adherent cells, as excessive washing could reduce the cell population available for downstream analysis. Cells were lysed in 200 µL of RIPA lysis buffer, followed by incubation on ice at 4°C for 5 minutes to ensure efficient cell lysis. After lysis, 500 µL of 1x PBS was added to detach the lysed material from the wells, and the wells were thoroughly washed to collect all possible cellular contents. The collected cell lysate was centrifuged at 4,000 rpm for 10 minutes at room temperature to separate the supernatant from any debris. This preparation ensured that the maximum amount of internalized DNA nanostructures could be retrieved for further analyses and selection processes. The supernatant was then transferred to a separate tube for purification of internalized structures.

#### 4.3.3 Cell lysate Processing

Cell lysate DNA was purified using AMPure XP paramagnetic beads (Beckman Coulter). The 700 µL sample was split into two tubes (350 µL per tube), and AMPure XP beads were added at a 1.8× ratio (630 µL per tube). Samples were mixed thoroughly and incubated for 10 min at room temperature to allow DNA binding. Following magnetic separation, the supernatant was removed, and the bead-bound DNA was washed twice with 500 µL of freshly prepared 75% ethanol. Beads were air-dried briefly, and DNA was eluted in nuclease-free water with a 10 min incubation before magnetic separation and transfer of the eluate to a fresh tube for further amplification and downstream processing

#### 4.3.4 PCR amplification of lysate

PCR reactions were set up using the conditions described in Section 4.1.1. Amplification was performed using DreamTaq 2× Master Mix (Thermo Fisher Scientific) in a total reaction volume of 1 mL, which was subsequently divided into ten 100 µL reactions, with template DNA at a final concentration of 1 ng/µL. Thermal cycling consisted of an initial denaturation at 95 °C for 2 min, followed by 15 cycles of denaturation at 95 °C for 30 s, annealing at 60 °C for 30 s, and extension at 72 °C for 30 s, with a final extension at 72 °C for 10 min.

The amplified products were purified using AMPure XP beads following the lysate purification protocol. A portion of the purified amplicons was used for nanopore sequencing, while the remaining sample was subjected to λ exonuclease digestion as described in Section 3.1.3 and subsequently refolded in 1× T4 ligase buffer.

### 4.4 Nanopore Sequencing

Amplicon libraries were prepared for Nanopore sequencing on the minion platform with flongle flow cells using the ONT Ligation Sequencing Kit (SQK-LSK114), following the manufacturer’s recommended workflow for the *Ligation Sequencing Amplicons V14* protocol.

#### 4.4.1 End Prep for nanopore sequencing

End Prep was performed to prepare the DNA ends for adapter ligation. Purified PCR amplicons were diluted in nuclease-free water to achieve 100 fmol input, assuming an average fragment length of 0.5 kb (32.5 ng DNA). DNA concentration was determined by Qubit, and the input volume was adjusted accordingly. End-prep reactions (30 µL total volume) were assembled as follows:

| Component | Volume ($\mu\text{L}$) |
| --- | --- |
| DNA amplicon | 24.5 |
| DCS | 0.5 |
| Ultra II End-prep Reaction Buffer | 3.5 |
| Ultra II End-prep Enzyme Mix | 1.5 |
| **Total** | **30.0** |

Reactions were mixed gently and incubated at 20 °C for 5 min followed by 65 °C for 5 min. DNA was purified using AMPure XP beads (30 µL), incubated for 5 min at room temperature, and magnetically separated. Beads were washed twice with freshly prepared 80% ethanol, briefly air-dried, and DNA was eluted in 30 µL nuclease-free water.

#### 4.4.2 Adapter ligation

Adapter ligation was performed to attach sequencing adapters to the DNA ends. To retain all DNA fragments, Short Fragment Buffer (SFB) was used during post-ligation clean-up. Ligation reactions (50 µL total volume) were assembled as follows:

| Component | Volume (μL) |
| --- | --- |
| End-prepped DNA | 30.0 |
| Ligation Buffer (LNB) | 12.5 |
| NEB Next Quick T4 DNA Ligase | 5.0 |
| Ligation Adapter (LA) | 2.5 |
| **Total** | **50.0** |

The reaction was incubated for 10 mins at room temperature. Post-ligation the sample was purified using AMPure XP beads (20 µL beads in 50 µL of reaction vol.) incubated on rotatory mixer for 5mins. The samples were briefly centrifuged and placed on a magnetic rack to remove supernatant, and the pelleted beads were washed twice with SFB, briefly air-dried, and eluted in 7 µL Elution Buffer. Library concentration was quantified using a Qubit fluorometer, and the final library was diluted to 5–10 fmol prior to sequencing.

#### 4.4.3 Flow cell check and sequencing

Flongle flow cells were inspected for active pores using MinKNOW software prior to sequencing. The DNA library was prepared at 5–10 fmol in 5 µL of Elution Buffer, corresponding to 3.25 ng of DNA for 10 fmol assuming an average amplicon length of 0.5 kb. Flush solution was prepared using 117 µL Flow Cell Flush (FCF) and 3 µL Flow Cell Tether (FCT) and used for priming the flow cell. The library sample was prepared as following:

| Component | Volume (μL) |
| --- | --- |
| Sequencing Buffer (SB) | 15 |
| Library Beads (LIB) | 10 |
| DNA library | 5 |
| **Total** | **30** |

The prepared library was loaded carefully to avoid air bubbles onto the flow cell, and sequencing was performed using super-accurate basecalling with a minimum read length threshold of 20 bp and POD5 output format.

### 4.5 Illumina Sequencing

We tested a total of five samples for Illumina sequencing which were: initial library, HEK293T-R5, HEK293T-R10, RAW264.7-R5 and RAW264.7-R10.

#### 4.5.1 Illumina Library Preparation

Libraries were prepared using a two-step PCR strategy to append Illumina-compatible adapters and unique dual-indices.

##### Primary Amplification (PCR-1)

The initial PCR was performed in a 25 µL reaction volume using 12.5 µL 2× KAPA HotStart ReadyMix (Roche), 1 µM each of custom primers containing Illumina-specific overhangs and library-specific regions, and 5–10 ng of template DNA. Cycling conditions consisted of an initial denaturation at 95°C for 3 min, followed by 8 cycles of 98°C for 20 s, 60°C for 15 s, and 72°C for 1 min, with a final extension at 72°C for 1 min. Primers used for this reaction were:

FWD Primer: 5’-ACACTCTTTCCCTACACGACGCTCTTCCGATCTCCAGCACGTGATATGTATATGT-3’
BWD Primer: 5’-GTGACTGGAGTTCAGACGTGTGCTCTTCCGATCTCCTACACGCGTACATGTATAGA-3’

##### Indexing and Adapter Ligation (PCR-2)

Secondary PCR amplification was used to incorporate P5/P7 adapters and unique indices. The 40 µL reaction comprised 20 µL 2× KAPA HiFi HotStart ReadyMix, 10 µL of purified PCR-1 product, and 1 µM indexed primers. The profile included an initial denaturation at 98°C for 2 min, followed by 8 cycles of 98°C for 20 s, 55°C for 30 s, and 72°C for 30 s, concluding with a 2 min extension at 72°C.Following each PCR step, products were purified using AMPure XP beads (Beckman Coulter) at a 1.8:1 volumetric ratio. Purification efficiency and the removal of primer dimers were confirmed via 2% agarose gel electrophoresis.

#### 4.5.2 Quantification and Equimolar Pooling

Library concentrations were determined via relative qPCR using SYBR Green chemistry, with the "initial library" sample serving as the reference standard. Primer efficiencies were derived from Ct values and maintained within an acceptable range of 90–110%. Based on these calculations, all the libraries were pooled in equimolar ratios to achieve a final concentration of ≈15 nM in a total volume of 54 µL.

### 4.6 Bioinformatics

#### 4.6.1 Sequencing data pre-processing for downstream analysis

Nanopore sequencing was performed on the MinION Mk1B device via the Flongle adapter using the MinKNOW software. For each run, sequencing was performed over 24 hours. Sequencing reads were pre-filtered, setting the minimal Q score to 10. Basecalling was performed using Dorado’s super-accurate basecalling. The fastQ files containing the passed and basecalled reads were concatenated for downstream processing.

The Illumina reads were split by index into five groups representing the five sequenced libraries. Illumina sequencing results in two paired end reads up to 300 nt long each, read from either end, before further processing these paired reads were fused using the tool Pear ^38^ into one single non-overlapping read.

#### 4.6.2 cONcat: Mapping of the structure genome to initial library fragments

To identify and decode the known fragments in each read i.e. structure genome, the concatenated fastQ files were run through cONcat, a software developed by Petri et al.,.^28^ Here, the concatenated fastQ files were given to the software together with the sequences of the fragments making up the components of the initial library (**Table S1**). The fragment sequences were mapped onto the reads using an edit-distance based recursive covering algorithm with an identity threshold of 0.75. If present, the software returns a set of best matched fragments above the identity threshold for each read, including the start and end positions of the fragments within the reads, the fragment name and edit distance of the mapping. This information was used to further decode general trends of fragment pairings, order, and frequency in the structure genomes throughout the selection process.

#### 4.6.3 UMI extraction and counting of barcodes

To further analyze the reads from the sequencing data and identify and count the prevalence of unique structure genomes, each read was characterized and sorted by its barcode, consisting of UMI1 and UMI2. To extract the UMIs from the sequencing data, concatenated fastQ files were processed using python, utilizing Bio.Align and Bio.SeqIO packages. To ensure that sequencing reads corresponded to true structure genomes, reads with a length below 80 nucleotides were discarded. The remaining reads were scanned for inclusion of the universal primer binding sites. Using the known primer sequences for the forward and reverse primer, primer binding sites were localized via pairwise local alignment within the first and last 90 nucleotides. Any read with a primer alignment score below xx threshold was discarded. Since the ONT Ligation Sequencing Kit (SQK-LSK114) results in reads with mixed read directions, backward reads were turned into their reverse complement strand, resulting in a pool of ‘pure’ forward reads. Via the forward primer alignment, UMI1 was defined to be the 10 nucleotides following the forward primer binding site. Similarly, UMI2 was defined as the 10 nucleotides prior to the backward primer alignment. UMI1 and UMI2 sequences were fused into a barcode, with each unique barcode denoting a unique structure genome. Reads were then sorted by barcode, and each barcode was counted to further analyze the enrichment of unique structure genomes.

#### 4.6.4 Detection and filtering of biological sequences in data

In order to detect data of structure genomes that appear to be chimeras with genetic DNA from the cells we aligned all reads to the GRCh38 human reference genome using minimap2. Reads that were mapped to the reference genome were filtered out and not used in selection of candidates from round 10 from HEK and RAW as well as control candidates.

#### 4.6.5 Secondary structure prediction of selected candidates

The secondary structures of selected DNA candidates were predicted using the Zuker mfold web server utilizing UNAfold software package configured for DNA folding^29^. The folding simulations were conducted under following conditions: 100 mM [Na⁺], 10 mM [Mg²⁺], and a temperature of 37 °C. The folding algorithm was configured with a percent suboptimality of 5%, ensuring that only structures within 5% of the minimum free energy (MFE) were considered. The upper bound on the number of computed foldings was restricted to 5 to prioritize the most thermodynamically probable architectures. A window parameter of 10 was applied to control the diversity of the generated structures, preventing the output of highly similar, redundant conformations.

The minimum Gibbs free energy (Δ*G*) of self-hybridization was calculated for each sequence to quantify the energetic favorability of the predicted folds. To facilitate a normalized comparison across candidates of varying lengths, the Stability Density was calculated as follows:

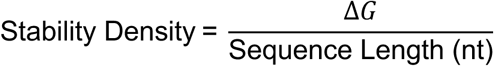

### 4.7 Candidate Synthesis and Verification

#### 4.7.1 Amplification from Purified Lysate

The initial candidate series (H5A–H5D) was amplified using purified lysate as the primary template at a concentration of 2ng/µL. Specific primer sets were designed for each candidate (Table S2), with the reverse primers conjugated to a Cy5 fluorophore to facilitate downstream cellular assays. To ensure high specificity and yield, gradient PCR was employed to determine the ideal annealing temperature based on the thermodynamic profiles of each primer pair. The reaction was performed in a 100µLvolume using the setup detailed in Table 1.

| Component | Volume (μl) |
| --- | --- |
| 2x DreamTaq Master Mix | 50 |
| Fwd primer (10 μM) | 2.5 |
| Bwd primer (10 μM) | 2.5 |
| Template (Purified Lysate) | 5 |
| H2O | 40 |
| Total Vol (ul) | 100 |

Thermocycling conditions:

- Initial Denaturation 95°C for 3 minutes
- 30 Cycles:
  - Denaturation: 95°C for 30 seconds
  - Annealing: 50°C for 30 seconds
  - Extension: 72°C for 30 seconds
- Final Extension: 72°C for 5 minutes

#### 4.7.2 Amplification from commercial template

The finalized candidates were synthesized from ANSA as a purified DNA fragments and were received as lyophilized material. These candidates were then resuspended in nuclease free water in the stock concentration of 100nM and that stock was further diluted to 10nM using nuclease free water. This template was then used for amplification using UMI specific primers. Here is the reaction setup for finalized candidates:

| Component | Volume (μl) |
| --- | --- |
| 2x DreamTaq Master Mix | 50 |
| Fwd primer (10 μM) | 1 |
| Bwd primer (10 μM) | 1 |
| Template (Purified Lysate) | 0.5 |
| H2O | 47.5 |
| Total Vol (ul) | 100 |

##### Thermal Cycling Conditions (Commercial)

- Initial Denaturation 95°C for 5 minutes
- 22 Cycles:
  - Denaturation: 95°C for 30 seconds
  - Annealing: 60°C for 30 seconds
  - Extension: 72°C for 30 seconds
- Final Extension: 72°C for 5 minutes

#### 4.7.3 Post amplification processing and verification

PCR amplification efficiency and product integrity were assessed by agarose gel electrophoresis. Amplification products were separated on a 2% (w/v) agarose gel prepared in 1× Tris–Borate–EDTA (TBE) buffer and stained with SYBR™ Safe DNA Gel Stain for nucleic acid visualization.

Gel imaging was performed using a Bio-Rad Gel Doc™ documentation system to verify the presence of both the DNA backbone and the fluorophore label. DNA bands were visualized using standard SYBR Safe acquisition settings. Detection of Cy5-labeled primers and their corresponding DNA products was achieved by operating the system in blot mode, with simultaneous signal acquisition in the AF488 (SYBR Safe) and Cy5 channels. Dual-channel imaging enabled confirmation of incorporation of Cy5 labelled primers in the amplified structures.

Following PCR, all candidates were purified using AmPure XP magnetic beads to remove unincorporated primers and reaction artifacts. To generate single-stranded DNA or specific structural intermediates, the purified products were subjected to lambda exonuclease digestion. Subsequently, the candidates were refolded in 1x T4 DNA ligase buffer to ensure proper structural assembly.

### 4.8 Flow cytometry

Cells were seeded in 12-well tissue culture plates one day prior to experimentation and maintained under standard culture conditions (37 °C, 5% CO₂) to ensure optimal adherence and recovery. On the day of stimulation, the culture medium was replaced with fresh serum-free medium containing DNA nanostructures at a final concentration of 25 nM and incubated for 1 h. Following incubation cells were washed with 1× PBS, detached via trypsinization, and transferred to U-bottom 96-well plates. Each experimental condition was performed in biological triplicates or quadruplets. To eliminate unbound nanostructures, cells were washed three times with 1× PBS by centrifugation at 800 rpm for 5 min.

The cell pellets were subsequently fixed with 2% paraformaldehyde (PFA) for 10–15 min at room temperature, washed twice with 1× PBS, and resuspended in FACS staining buffer (1× PBS supplemented with serum). Samples were stored at 4 °C protected from light and analyzed the same day. Untreated cells were included as negative controls to determine baseline fluorescence.

Flow cytometry data were acquired using a BD FACSCanto II system equipped with BD FACSDiva software, with 10,000 events recorded per sample. Data analysis and visualization were performed using FlowJo software employing conventional gating strategies.

### 4.9 Confocal Microscopy

Fixed cells were imaged using a Leica Stellaris X confocal scanning laser microscope (LIA). Samples were visualized using a 63× oil-immersion objective or a 40× glycerol objective, as specified in the figure legends. Fluorophores were excited using the following laser lines: Hoechst (405 nm), LysoTracker Green (488 nm), and Cy5/TYE-labeled DNA candidates (633 nm). To ensure imaging accuracy and eliminate spectral crosstalk, images were acquired using sequential scanning mode. For each sample, 3–4 Z-stacks were captured to assess and ensure representative sampling. Representative images displayed in the figures are projections generated from several fused optical sections from the middle of the Z-stack to clearly illustrate internal localization.

Image processing was performed using Fiji (ImageJ, NIH). Processing was limited to background subtraction and brightness/contrast adjustments applied uniformly across all images within a comparison set to ensure representative visualization of signal intensities.

## Statistics

Statistical analyses were performed using GraphPad Prism 10. Data are presented as mean±standard deviation (SD). For quantitative comparisons of flow cytometry data statistical significance was determined using two-tailed unpaired t-tests. A p-value ≤0.05 was defined as the threshold for statistical significance. Regarding experimental design, procedures were not randomized, and investigators were not blinded to group allocation during data collection or outcome assessment. All error bars presented in figures represent the SD of at least 3 independent biological replicates or as mentioned in the figure.

## Acknowledgements

We thank the Birgitta Henriques-Normark and group for helping us to use their cell culture facility. We thank Marie-Steph and Rebecca Dookie for providing the cell lines. We are grateful to the National Genomics Infrastructure (NGI) at SciLifeLab for their assistance with Illumina sequencing. We also thank the Biomedicum Imaging Core (BIC) and the Biomedicum Flow Cytometry Core Facility (BFC) at Karolinska Institutet for help with confocal microscopy and flow cytometry, respectively.

## Authors Contribution

AR: Investigation, writing—original draft preparation, review, and editing.

LS: Data analysis, review, and editing.

JP: AFM analysis and review.

SL: Illumina sequencing.

EB: Conceptualization, writing, editing, data analysis, and funding acquisition.

