## Supplementary tables and figures for "Evolutionary selection of DNA nanostructures for cellular uptake"

#### Supplementary table 1: Sequence information of individual fragments used for library preparation

| Name | Sequence |
|---|---|
| 3WJ Strand1 | /5Phos/ACTCTCTGTATGTAGCATACGCGCACG |
| 3WJ Strand2 | /5Phos/AGTCGTGCGCGTATGCGTAGCTGCGAG |
| 3WJ Strand3 | /5Phos/ACTCTCGCAGCTACGCTACATACAGAG |
| 3WJ unpairedT Strand1 | /5Phos/ACTCTGCACGTGTGGTCGTGTATCAGCG |
| 3WJ unpairedT Strand2 | /5Phos/AGTCGCTGATACACGTCCGATGATACGG |
| 3WJ unpairedT Strand3 | /5Phos/ACTCCGTATCATCGGTCCACACGTGCAG |
| 4WJ Strand1 | /5Phos/ACTCGCTATCTACGGCCGCATGATACG |
| 4WJ Strand2 | /5Phos/AGTCGTATCATGCGGCGCATATCATGG |
| 4WJ Strand3 | /5Phos/ACTCCATGATATGCGCCTGTGCATGAG |
| 4WJ Strand4 | /5Phos/ACTCTCATGCACAGGCCGTAGATAGCG |
| KL-10nt Strand1 | /5Phos/AGTCCATCATGTGCAGGCATGCGCTACCCTGCACATGATGG |
| KL-10nt Strand2 | /5Phos/AGTCTACATGATACGTGGTAGCGCATGCACGTATCATGTAG |
| KL-12nt-Strand1 | /5Phos/AGTCTATCTGTACATGGTGCTATGATATACCATGTACAGATAG |
| KL-12nt-Strand2 | /5Phos/AGTCTGTATCACACGCGTGCTATGATATACGCGTGTGATACAG |
| KL-14nt-Strand1 | /5Phos/AGTCCTATACACAGCAGTGTGCATATGTATGCTGCTGTGTATAGG |
| KL-14nt-Strand2 | /5Phos/AGTCTATACACACGCTGCATACATATGCACACAGCGTGTGTATAG |
| KL-16nt-Strand1 | /5Phos/AGTCACATATATCTGCGCGTAGATAGCTGCTGACGCAGATATATGTG |
| KL-16nt-Strand2 | /5Phos/AGTCATCATATACGCGGTCAGCAGCTATCTACGCCGCGTATATGATG |
| KL-18nt-Strand1 | /5Phos/AGTCACGTAGCGCACAGGATACAGCAGCTATGTAGCTGTGCGCTACGTG |
| KL-18nt-Strand2 | /5Phos/AGTCTGTGCACAGATAGCTACATAGCTGCTGTATCCTATCTGTGCACAG |
| KL-20nt-Strand1 | /5Phos/AGTCTATCGATAGCTGGTCTACAGATACATGCGCGTACCAGCTATCGATAG |
| KL-20nt-Strand2 | /5Phos/AGTCCTATACAGATAGGTACGCGCATGTATCTGTAGACCTATCTGTATAGG |
| Amplification Loop | /5Phos/ ACTCCATACGTATCTANNNNNNNNNNACATATACATATCACGTGCTGGCCTACACGCGTACATGTATAGANNNNNNNNNNTAGATACGTATGG |
| Hairpin 10nt | /5Phos/AGTCGCACACGATACAGACACACATGACTGTATCGTGTGC |
| Hairpin 12nt | /5Phos/AGTCCGCATCATATGTACGCATCAGATCATATGATGCGG |
| Hairpin 14nt | /5Phos/AGTCATATCGATATGTGACATATCATCGCGTCACATATCGATATG |
| Hairpin 16nt | /5Phos/AGTCCATCTGTGTGCGGTGCGCATGATATAGATCCGCACACAGATGG |
| Hairpin 18nt | /5Phos/AGTCCTATCGATATATGCAGATATCGTATCGCAGCCATATATCGATAGG |
| Linear 10nt Strand1 | /5Phos/AGTCACGTGATACGG |
| Linear 10nt Strand2 | /5Phos/ACTCCGTATCACGTG |
| Linear 11nt Strand1 | /5Phos/AGTCATACGCGTATAG |
| Linear 11nt Strand2 | /5Phos/ACTCTATACGCGTATG |
| Linear 12nt Strand1 | /5Phos/AGTCCGCAGCGTACGAG |
| Linear 12nt Strand2 | /5Phos/ACTCTCGTACGCTGCGG |
| Linear 13nt Strand1 | /5Phos/AGTCCATCATCACATATG |
| Linear 13nt Strand2 | /5Phos/ACTCATATGTGATGATGG |
| Linear 14nt Strand1 | /5Phos/AGTCATCAGCTGCATGTGG |
| Linear 14nt Strand2 | /5Phos/ACTCCACATGCAGCTGATG |
| Linear 15nt Strand1 | /5Phos/AGTCCTGCATCTATCGCTAG |

| | |
|---|---|
| Linear 15nt Strand2 | /5Phos/ACTCTAGCGATAGATGCAGG |
| Linear 16nt Strand1 | /5Phos/AGTCGATATATAGCGCTGCGG |
| Linear 16nt Strand2 | /5Phos/ACTCCGCAGCGCTATATATCG |
| Linear 17nt Strand1 | /5Phos/AGTCTATCTGCTAGCTGTGTGG |
| Linear 17nt Strand2 | /5Phos/ACTCCACACAGCTAGCAGATAG |
| Linear 18nt Strand1 | /5Phos/AGTCTAGCTACGCGTATATCGTG |
| Linear 18nt Strand2 | /5Phos/ACTCACGATATACGCGTAGCTAG |
| Linear 19nt Strand1 | /5Phos/AGTCCGCTGATATCTACACACGCG |
| Linear 19nt Strand2 | /5Phos/ACTCGCGTGTGTAGATATCAGCGG |
| Linear 20nt Strand1 | /5Phos/AGTCTCTATCGCGCAGCTATCACAG |
| Linear 20nt Strand2 | /5Phos/ACTCTGTGATAGCTGCGCGATAGAG |
| Linear 1nt bulge Strand1 | /5Phos/AGTCTATGTGCGATGCTATCATGTG |
| Linear 1nt bulge Strand2 | /5Phos/ACTCACATGATAGCACGCACATAG |
| Linear 2nt bulge Strand1 | /5Phos/AGTCATCATGATACGTGTACGCTAG |
| Linear 2nt bulge Strand2 | /5Phos/ACTCTAGCGTACACGTCATGATG |
| Linear 3nt bulge Strand1 | /5Phos/AGTCCGATATATACGTAGCACGCAG |
| Linear 3nt bulge Strand2 | /5Phos/ACTCTGCGTGCTACTATATCGG |
| Linear 4nt bulge Strand1 | /5Phos/AGTCCATCACATACAGCGTAGCTAG |
| Linear 4nt bulge Strand2 | /5Phos/ACTCTAGCTACGCTGTGATGG |
| Linear 5nt bulge Strand1 | /5Phos/AGTCATATCACATATCGCACGTAGG |
| Linear 5nt bulge Strand2 | /5Phos/ACTCCTACGTGTGTGATATG |
| Forward Primer | /5Phos/CCATATGTGTACACGTGTGATA |
| Reverse Primer | CCTGCGCAGCTATATCTATCAG |

**Supplementary table 2: Cluster Primer Sequences**

| Name | Sequence |
|---|---|
| H5A_FWD | /5Phos/ATGTATATGTTAACGAAGAA |
| H5A_BWD | /5Cy5/CATGTATAGACAAGCGATAA |
| H5B_FWD | /5Phos/ATGTATATGTGGACTTCAGG |
| H5B_BWD | /5Cy5/CATGTATAGACGCCGATTGG |
| H5C_FWD | /5Phos/ATGTATATGTATCTCCCAGA |
| H5C_BWD | /5Cy5/CATGTATAGATATTGGCATC |
| H5D_FWD | /5Phos/ATGTATATGTTGACCTTACC |
| H5D_BWD | /5Cy5/CATGTATAGAGGAGTGAGAG |
| STD_FWD | /5Phos/CCAGCACGTGATATGTATATGT |
| STD_REW | /5Cy5/CCTACACGCGTACATGTATAGA |

**Supplementary table 3: Finalized candidates and their corresponding sequence information**

| Name | Sequence |
|---|---|
| RAW10A | CCATATGTGTACACGTGTGATA-<br>ACGCGTGTAGGCCAGCACGTGATATGTATATGCCTCCACATGCAGCTGATGACT<br>CTGTGATAGCTGCGCGATAGAGACTCTAGCTACGCTGTATGTGATGGACTCCGC<br>ACAGCTAGCAGATAGACTCATATGTGATGATGGACTCTAGCTACGCTGTATGTGA<br>TGGACTCCATGATATGCGCCGCATGATACGACTCCATACGTATCTAGCTCCCCC<br>CC-CTGATAGATATAGCTGCGGAGG |
| RAW10B | CCATATGTGTACACGTGTGATA-<br>ACGCGTGTAGGCCGGCACGTGATATGTATATGTTATGTTTACCTAGATACGTATG<br>GAGTCCATCACAGCGTAGCTAGAGTCTATGTGCGTGCTATCATGTGAGTCCTATA<br>TAGGTCTACAGATACATGCGCGTACCTATCTGTATAGGACTCCTACGTGCTATAC<br>GTGATATGACTCCGTATCATCGGACGTGTATCAGCGACTCCTACGTGCGATATGT<br>GATATGACTCCATACGTATCTACCCCGCCCCC-CTGATAGATATAGCTGCGGAGG |
| RAW10C | CCATATGTGTACACGTGTGATA-<br>ACGCGTGTAGGCCAGCACGCGATATGTATATGTCACGCTCGTCTAGGTACGTAT<br>GATCTGATGCGTACATATGATGCGGACTCCACACAGCTAGCAGACAGACTCCAC<br>ACAGCTAGCAGATAGACTCGCGTGTGTAGATATCAGCGGACTCTCGCAGCTACG<br>TATACGCGCACGACTCCGTATCACGTGACTCACGATATACGCGTAGCTAGACTC<br>CACATGCAGCTGATGACTCCTACGTGCGATATGTGATATGACTCCATACGTATCT<br>ACCCTACCTCTTCTATATATGTACGCGTGTAGGCCAGCACGTGATATGTATACGT<br>TGCTTCTTCCTAGATACGTATGGAA-CTGATAGATATAGCTGCGGAGG |
| RAW10D | CCATATGTGTACACGTGTGATA-<br>ACGCGTGTAGGGCAGCACGTGATATGTATATGTGACAGCACCATAGATACGTAT<br>GGAGTCCTATCGATATATGGCTGCGATACGATATCTGCATATATCGATAGGACTC<br>TAGCGTACACGATCATGATGACTCCACATGCAGCTGATGACTCCACATGCAGCT<br>GATGACTCCACATGCAGCTGATGACTCCATACGTATCTACCCCTCTCCG-<br>CTGATAGATATAGCTGCGGAGG |
| H10A- | CCATATGTGTACACGTGTGATA-<br>GGTTGCCAAATAGATACGTATGGAGTCACGTGATACGGAGTCTGTGCACAGATA<br>GGATACAGCAGCTAGGACATAAGCTCTATACATGTACGTGTGTAGGCCAGCACG<br>TGATATGTATATGTGGTTGCCAAATAGATACGTATGGAGTCACGTGATACGGAGT<br>CTGTGCACAGATAGGATACAGCAGCTAGGACATAAGC-<br>CTGATAGATATAGCTGCGGAGG |
| H10B | CCATATGTGTACACGTGTGATA-<br>GAGGTGACGGGTATCATCGGACGTGTATCAGCGACTCCGTATTATCGGACGTGT<br>ATCAGCGACTCCATACGTATCTACAGTGAGTGGTCTATACATGTACGCGTGTAGG<br>CCAGCACGTGATATGTATATGTGAGGCGACGGGTATCATCGGACGTGTATCAGC<br>GACTCCGTATCATCGGACGCGTATCAGCGACTCCATACGTATCTACAGTGAGTG<br>G-CTGATAGATATAGCTGCGGAGG |
| H10C | CCATATGTGTACACGTGTGATACAAACAAACTTAAATACGTACAAATAAACTCCAC<br>ATGCAGCTAATAACTCACAACATACGCGTAGCTAAACTCCATACGTATCTAACAG<br>GAACGCTCTATACATGTACGCCGTGGAGGCCAGCACGTGATATGTATGTGTCAA<br>ACAAACTTAAATACGTACAAATAGACTCCACATGCAGCTGATGACTCACGATATA<br>CGCGTAGCTAGACTCCATACGTATCCAACAGGAACGC-<br>CTGATAGATATAGCTGCGGAGG |
| H10D | CCATATGTGTACACGTGTGATAACGCGTGTAGGCCAGCACGTGATATGTATATGT<br>TAAAGGGCATTAGATACGTATGGAGTCCATCACAGCGTAGCTAGAGTCATCAGCT<br>GCATGTGGAGTCCATCATCACATATGAGTCATCATATACGCGGCGTAGATAGCC<br>GCTGACCGCGTATATGATGACTCTATACGCGTATGACTCCTATGTGCGATATGTG<br>ATATGACTCCATACGTATCTACCCCCCCCC-CTGATAGATATAGCTGCGGAGG |
| Crude Control C | CCATATGTGTACACGTGTGATA-<br>CCGCATTGCTTAGATACGTATGGAGTCATCATGACGTGTACGCTAGAGTCTATAC<br>ACACGCTGTGTGCATATGTATGCAGCGTGTGTATAGACTCTAGCGTACACGTATC<br>ATGATGACTCCATACGTATCTAGACAGTGCAC-CTGATAGATATAGCTGCGCAGG |

| | |
| --- | --- |
| | CCATATGTGTACACGTGTGATACCCACGAAAATAGATACGTATGGAGTCATCTAT |
| | ACATGTACGCGTGTAGGCCAGCACGTGATATGTATATGTTCATTAAGATCTAGAT |
| | ACGTATGGAGTCACATATATCTGCGTCAGCAGCTATCTACGCGCAGATATATGTG |
| Random-1 | ACTCCATACGTATCTAATCTCCCGCTCTGATAGATATAGCTGCGCAGG |
| | CCATATGTGTACACGTGTGATAACGCGTGTAGGCCAGCACGTGATATGTATATGT |
| | TAGACATACCTAGATACGTATGGAGTCCATCATCACATATGAGTCCGATATAGTA |
| | GCACGCAGAGTCCTATACACAGCAGCATACATATGCACACTGCTGTGTATAGGA |
| | CTCTGCGTGCTACGTATATATCGGACTCATATGTGATGATGGACTCCATACGTAT |
| Random-4 | CTACCGGCCGCTCCTGATAGATATAGCTGCGCAGG |
| | CCATATGTGTACACGTGTGATATTGCTTACATTAGATACGTATGGAGTCCTATACA |
| | GATAGGTCTACAGATACATGCGCGTACCTATCTGTATAGGACTCCATACGTATCT |
| | ATATGGTGCCTTCTATACATGTACGCGTGTAGGCCAGCACGTGATATGTATATGT |
| | TTGCTTACATTAGATACGTATGGAGTCCTATACAGATAGGTCTACAGATACATGC |
| | GCGTACCTATCTGTATAGGACCCCATACGTATCTATATGGTGCCTCTGATAGATA |
| Random-6 | TAGCTGCGCAGG |
| | CCATATGTGTACACGTGTGATAGGACTTCAGGAGTGGAACCACAGACCTTCACA |
| | GTGAGTGTTACAGCTCTTAAAGGTGGCGCGTCCGGAGTTGTTTGTTCCTCTCGG |
| | TGGGTTTGTGGTCTCACTGACTTCAGGAATGGAGTCTAGACCCTCGCAGTGAGT |
| | GTTACAGCTCATAAAAGTAGTGTGGACCCAAAGAGTGACCATCAGCAAGATTTAT |
| | TGTGAAGAGCGAAAGAACAAAGCTTCCACAGTATGGAAGGGGACCCACAGCTAG |
| | CGGATAGACTCCAGGACGCACTCACCAATTGGCGCTGATAGATATAGCTGCGCA |
| H5B | GG |
| | CCATATGTGTACACGTGTGATATGACCTTACCATACCTCATATGTTCAATTAACAC |
| | ACTATATTTTAAAACAGACTCAGATCTTGAAAACAAGATATAAGGTCCAATCCCTA |
| | TACATGGTGCGTAAGGTGGCTTTGGGGAATACAGTAGAAACCTAGTTTTCAGCAT |
| | GGCTCTCAGAAAACACCATCCCTCCTGCCTGTGGTAAGTATTCGTAGTAAGAGCT |
| | GAAACGGTGGGGTCAAGAGAACATCACTTCGGGTGAGAAAGTTTGTTTCATACTT |
| | TATTTCTCACAATTCTCGCCGTCAAGAGAGGAGTTTGATTCATCAACTTTGGGAT |
| | GCATTAGAATAACCCGTGGGATGGATTAAATAGATTCCATGGCCCTGCTCCTGAG |
| | AGATCCTTATTCAGTGGATGTGACGTGCGACTTAACTATCACTTAGCTTAATTTAC |
| | TTCACAGTCAGTGTACATATGCACACTGCTGTGTATAGAACTCCATACGTATCTA |
| H5D | CTCACTCCCTGATAGATATAGCTGCGCAGG |

### Supplementary table 4: UMI counts of candidates

Fraction of reads in each sequencing dataset (not filtered to remove biological data) that contains the fused UMI of the candidates used in this study. Pairwise alignment was used to also find UMI's that may contain read errors (primarily in ONT data).

| **Candidate Name** | | | | | | | | | | | | | |
|---|---|---|---|---|---|---|---|---|---|---|---|---|---|
| Cell lines | N/A | HEK293T | HEK293T | HEK293T | HEK293T | RAW264,7 | RAW264,7 | RAW264,7 | RAW264,7 | N/A | HEK293T | HEK293T | RAW264,7 | RAW264,7 |
| Round | 0 | 4 | 6 | 8 | 10 | 4 | 6 | 8 | 10 | 0 | 5 | 10 | 5 | 10 |
| Sequencing method | ONT | ONT | ONT | ONT | ONT | ONT | ONT | ONT | ONT | Illumina | Illumina | Illumina | Illumina | Illumina |
| H5A | 0,00E+00 | 8,98E-05 | 1,94E-01 | 4,01E-01 | 4,67E-01 | 1,65E-03 | 9,31E-03 | 5,26E-02 | 8,70E-03 | 5,78E-05 | 1,19E-06 | 2,17E-01 | 7,32E-04 | 3,09E-03 |
| H5B | 0,00E+00 | 1,26E-05 | 1,78E-03 | 1,82E-03 | 9,87E-04 | 0,00E+00 | 0,00E+00 | 1,85E-05 | 1,44E-05 | 2,75E-07 | 0,00E+00 | 8,98E-05 | 1,85E-07 | 2,36E-06 |
| H5C | 0,00E+00 | 3,59E-06 | 1,77E-03 | 3,65E-03 | 5,65E-03 | 1,91E-06 | 0,00E+00 | 4,63E-06 | 0,00E+00 | 1,37E-07 | 0,00E+00 | 1,96E-03 | 3,69E-07 | 0,00E+00 |
| H5D | 0,00E+00 | 1,80E-06 | 1,85E-04 | 5,16E-04 | 4,62E-04 | 0,00E+00 | 0,00E+00 | 0,00E+00 | 1,20E-06 | 4,12E-07 | 4,76E-07 | 7,00E-05 | 1,85E-07 | 3,37E-07 |
| H10A | 0,00E+00 | 1,80E-06 | 4,04E-05 | 9,66E-05 | 1,92E-04 | 0,00E+00 | 0,00E+00 | 0,00E+00 | 1,20E-06 | 1,37E-07 | 2,38E-07 | 6,77E-04 | 1,85E-07 | 1,35E-06 |
| H10B | 0,00E+00 | 1,80E-06 | 9,81E-05 | 1,53E-04 | 1,18E-04 | 0,00E+00 | 0,00E+00 | 0,00E+00 | 0,00E+00 | 1,37E-07 | 4,76E-07 | 4,12E-04 | 1,85E-07 | 1,01E-06 |
| H10C | 0,00E+00 | 0,00E+00 | 6,06E-05 | 6,18E-05 | 6,93E-05 | 0,00E+00 | 0,00E+00 | 0,00E+00 | 0,00E+00 | 2,75E-07 | 0,00E+00 | 1,10E-04 | 0,00E+00 | 0,00E+00 |
| H10D | 0,00E+00 | 1,39E-03 | 3,75E-03 | 3,83E-03 | 3,16E-03 | 5,98E-03 | 7,70E-03 | 8,93E-03 | 1,18E-02 | 2,95E-05 | 1,27E-04 | 5,36E-03 | 7,28E-03 | 1,35E-02 |
| R10A | 0,00E+00 | 1,03E-03 | 2,88E-03 | 3,19E-03 | 2,61E-03 | 4,95E-03 | 6,11E-03 | 7,31E-03 | 9,35E-03 | 2,25E-05 | 1,11E-04 | 4,54E-03 | 6,28E-03 | 1,16E-02 |
| R10B | 0,00E+00 | 1,36E-03 | 3,41E-03 | 3,55E-03 | 2,96E-03 | 6,20E-03 | 8,09E-03 | 9,33E-03 | 1,18E-02 | 3,23E-05 | 1,38E-04 | 5,21E-03 | 8,07E-03 | 1,42E-02 |
| R10C | 0,00E+00 | 3,82E-04 | 8,89E-04 | 5,99E-04 | 3,87E-04 | 1,45E-03 | 2,22E-03 | 2,40E-03 | 3,22E-03 | 6,45E-06 | 9,16E-04 | 9,03E-04 | 3,48E-03 | 6,19E-03 |
| R10D | 0,00E+00 | 8,74E-04 | 2,02E-03 | 1,80E-03 | 1,43E-03 | 4,06E-03 | 4,93E-03 | 5,66E-03 | 7,25E-03 | 1,52E-05 | 8,85E-05 | 2,44E-03 | 4,96E-03 | 8,73E-03 |
| Random-1 | 0,00E+00 | 0,00E+00 | 0,00E+00 | 0,00E+00 | 0,00E+00 | 1,91E-06 | 0,00E+00 | 0,00E+00 | 3,61E-06 | 5,49E-07 | 0,00E+00 | 0,00E+00 | 0,00E+00 | 5,05E-07 |
| Random-2 | 0,00E+00 | 0,00E+00 | 0,00E+00 | 0,00E+00 | 0,00E+00 | 0,00E+00 | 0,00E+00 | 0,00E+00 | 0,00E+00 | 1,37E-07 | 0,00E+00 | 0,00E+00 | 0,00E+00 | 0,00E+00 |
| Random-3 | 0,00E+00 | 0,00E+00 | 0,00E+00 | 0,00E+00 | 0,00E+00 | 0,00E+00 | 0,00E+00 | 0,00E+00 | 0,00E+00 | 1,37E-07 | 2,38E-07 | 0,00E+00 | 0,00E+00 | 0,00E+00 |
| Random-4 | 0,00E+00 | 0,00E+00 | 0,00E+00 | 0,00E+00 | 0,00E+00 | 0,00E+00 | 0,00E+00 | 0,00E+00 | 0,00E+00 | 9,61E-07 | 4,76E-07 | 0,00E+00 | 3,69E-07 | 0,00E+00 |
| Random-5 | 0,00E+00 | 0,00E+00 | 0,00E+00 | 0,00E+00 | 0,00E+00 | 0,00E+00 | 0,00E+00 | 0,00E+00 | 0,00E+00 | 4,12E-07 | 0,00E+00 | 0,00E+00 | 0,00E+00 | 0,00E+00 |
| Random-6 | 0,00E+00 | 0,00E+00 | 0,00E+00 | 0,00E+00 | 0,00E+00 | 0,00E+00 | 0,00E+00 | 0,00E+00 | 0,00E+00 | 1,37E-07 | 0,00E+00 | 0,00E+00 | 0,00E+00 | 0,00E+00 |
| Random-7 | 0,00E+00 | 0,00E+00 | 0,00E+00 | 0,00E+00 | 0,00E+00 | 0,00E+00 | 0,00E+00 | 0,00E+00 | 0,00E+00 | 1,37E-07 | 0,00E+00 | 0,00E+00 | 0,00E+00 | 0,00E+00 |
| Random-8 | 0,00E+00 | 0,00E+00 | 0,00E+00 | 0,00E+00 | 0,00E+00 | 0,00E+00 | 0,00E+00 | 0,00E+00 | 0,00E+00 | 2,75E-07 | 0,00E+00 | 0,00E+00 | 0,00E+00 | 0,00E+00 |
| Random-9 | 0,00E+00 | 0,00E+00 | 0,00E+00 | 0,00E+00 | 0,00E+00 | 0,00E+00 | 0,00E+00 | 0,00E+00 | 0,00E+00 | 5,49E-07 | 0,00E+00 | 0,00E+00 | 0,00E+00 | 0,00E+00 |
| Crude Crontrol A | 0,00E+00 | 0,00E+00 | 0,00E+00 | 0,00E+00 | 0,00E+00 | 0,00E+00 | 0,00E+00 | 0,00E+00 | 0,00E+00 | 1,37E-07 | 0,00E+00 | 0,00E+00 | 0,00E+00 | 0,00E+00 |
| Crude Crontrol B | 0,00E+00 | 0,00E+00 | 0,00E+00 | 0,00E+00 | 0,00E+00 | 0,00E+00 | 0,00E+00 | 0,00E+00 | 0,00E+00 | 1,37E-07 | 0,00E+00 | 0,00E+00 | 0,00E+00 | 0,00E+00 |
| Crude Crontrol C | 0,00E+00 | 0,00E+00 | 0,00E+00 | 0,00E+00 | 0,00E+00 | 0,00E+00 | 0,00E+00 | 0,00E+00 | 0,00E+00 | 1,37E-07 | 0,00E+00 | 0,00E+00 | 0,00E+00 | 3,37E-07 |
| Crude Crontrol D | 0,00E+00 | 0,00E+00 | 0,00E+00 | 0,00E+00 | 0,00E+00 | 0,00E+00 | 0,00E+00 | 0,00E+00 | 0,00E+00 | 1,37E-07 | 0,00E+00 | 0,00E+00 | 0,00E+00 | 0,00E+00 |

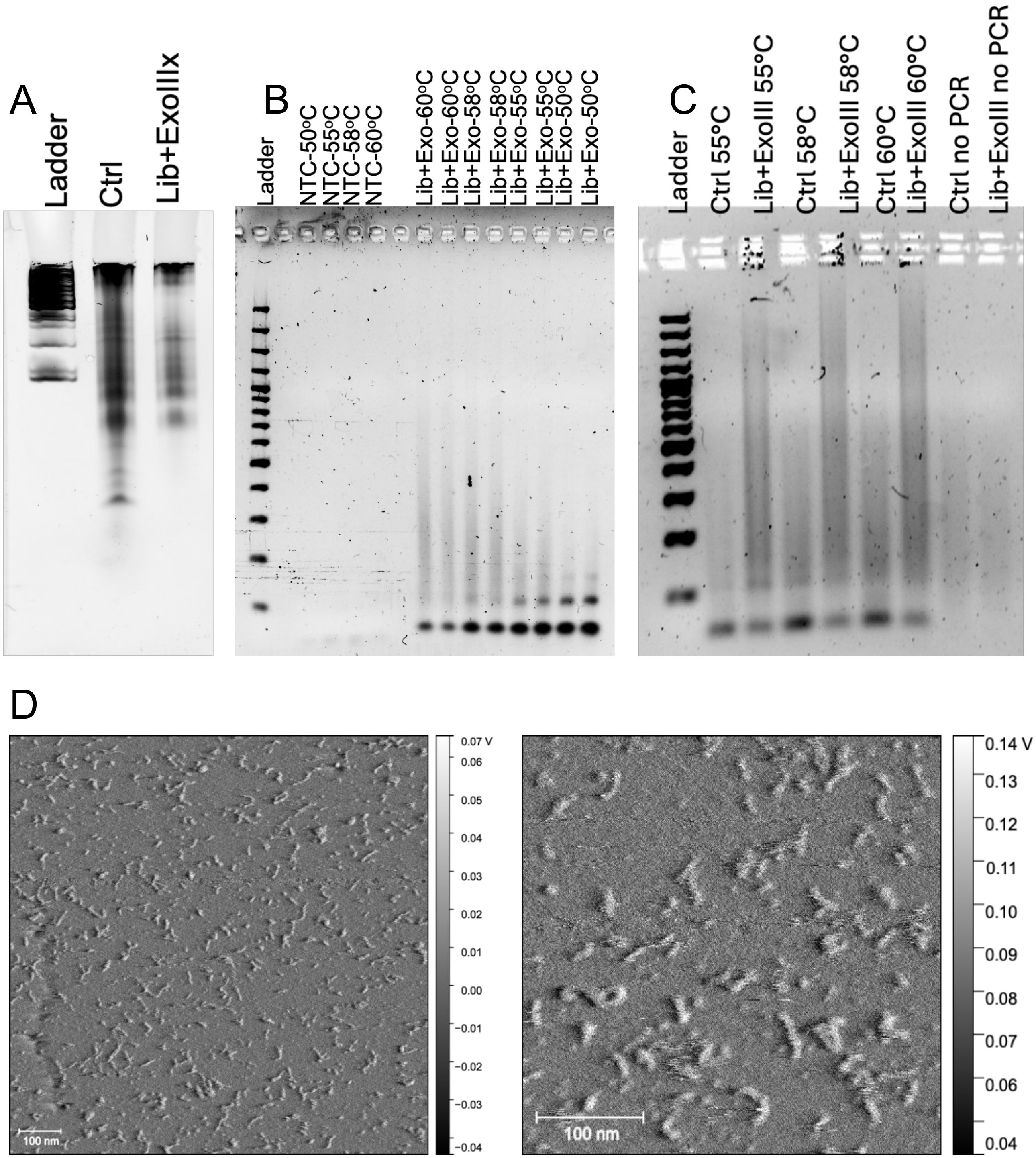


**Supplementary figure 1:** ***Preparation of DNA Library.*** (A) 10% urea PAGE stained with SYBR Gold, showing annealed and ligated fragments forming the library. "Ctrl" denotes the control, and "Lib+Exo" is the control library treated with exonuclease digestion, which removes partially unfolded or smaller fragments. (B) 2% agarose gel stained with SYBR Gold showing PCR amplification of the control library and Lib+Exo at different annealing temperatures. (C) 2% agarose gel stained with SYBR Gold showing lambda exonuclease digestion of the PCR-amplified and purified library. (D) AFM images of the amplified and purified DNA library. Scale bar: 100 nm.

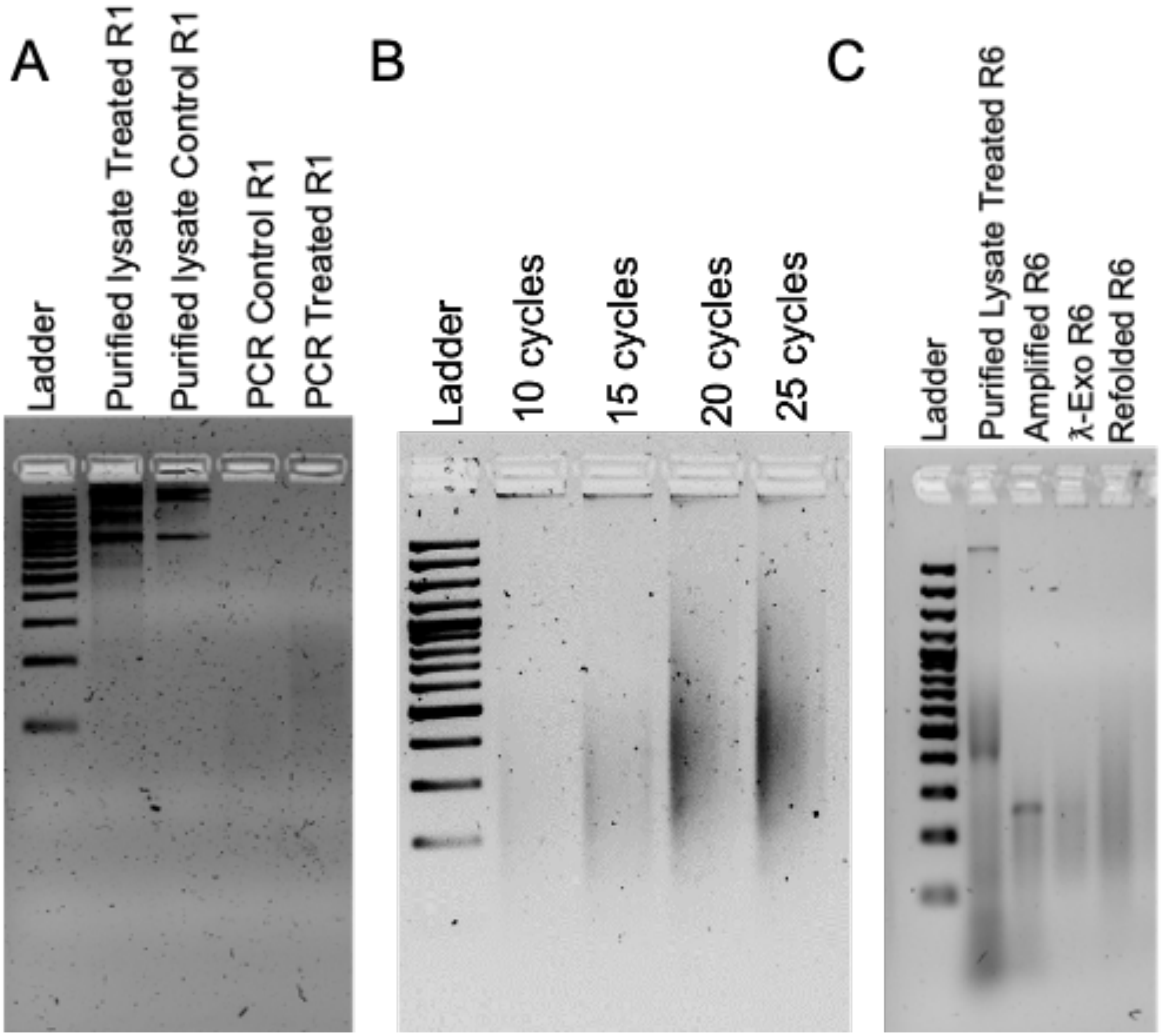


**Supplementary figure 2:** ***Cell Lysate Optimization for Subsequent Selection.*** (A) 2% agarose gel images showing purification of treated and non-treated (control) lysates and their PCR amplification. (B) SYBR Safe–stained 2% agarose gel showing PCR efficiency at different amplification cycles. (C) Representative SYBR Safe–stained 2% agarose gel showing purified Round-6 lysate, its amplification, λ-exonuclease digestion, and refolding for the next round of cell stimulation.

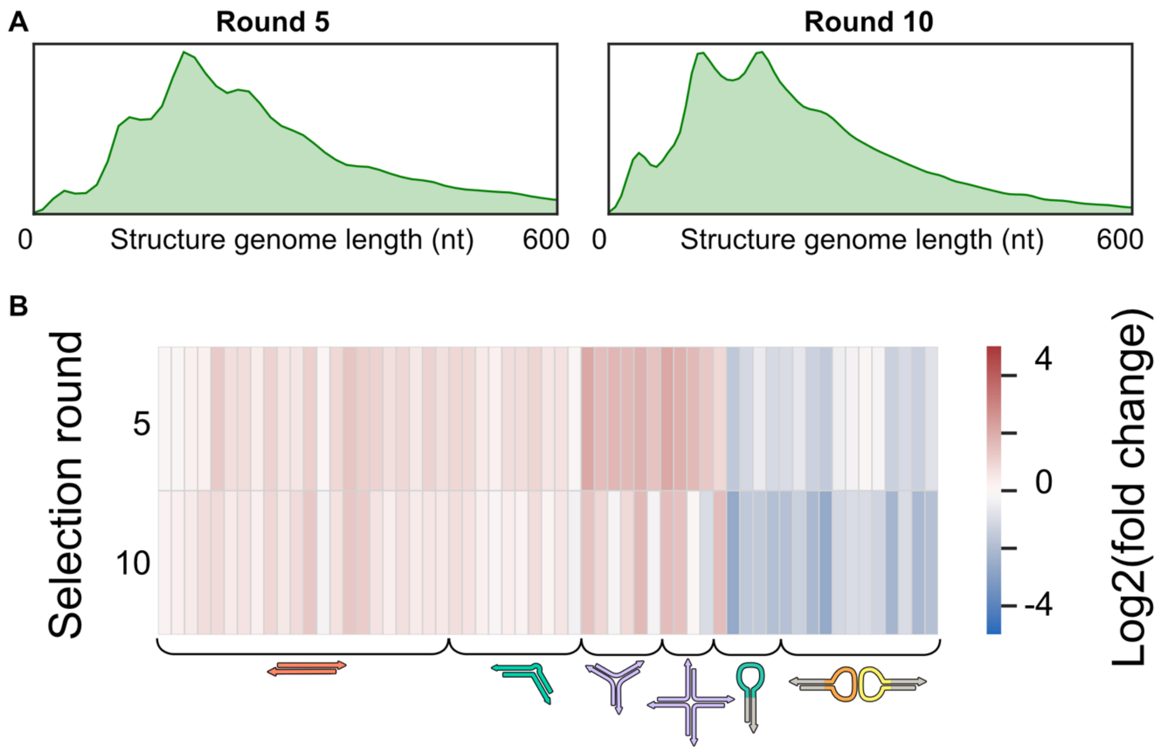


**Supplementary figure 3:** ***Empty selection and sequencing control.*** The Lib+ExoIII sample was amplified for 15 PCR cycles and purified using AMPure beads. This amplification and purification step was repeated for ten rounds without λ exonuclease digestion or refolding to assess amplification bias. **(A)** The initial DNA nanostructure library was PCR amplified 10 times to undertint the amplification preferences. The resulting products after 5 and 10 rounds of amplification was sequenced with Oxford nanopore. A) Distribution of genome lengths after 5 or 10 rounds of amplification. B) Structure fragment preference after 5 or 10 rounds of amplification compared to the crude library.

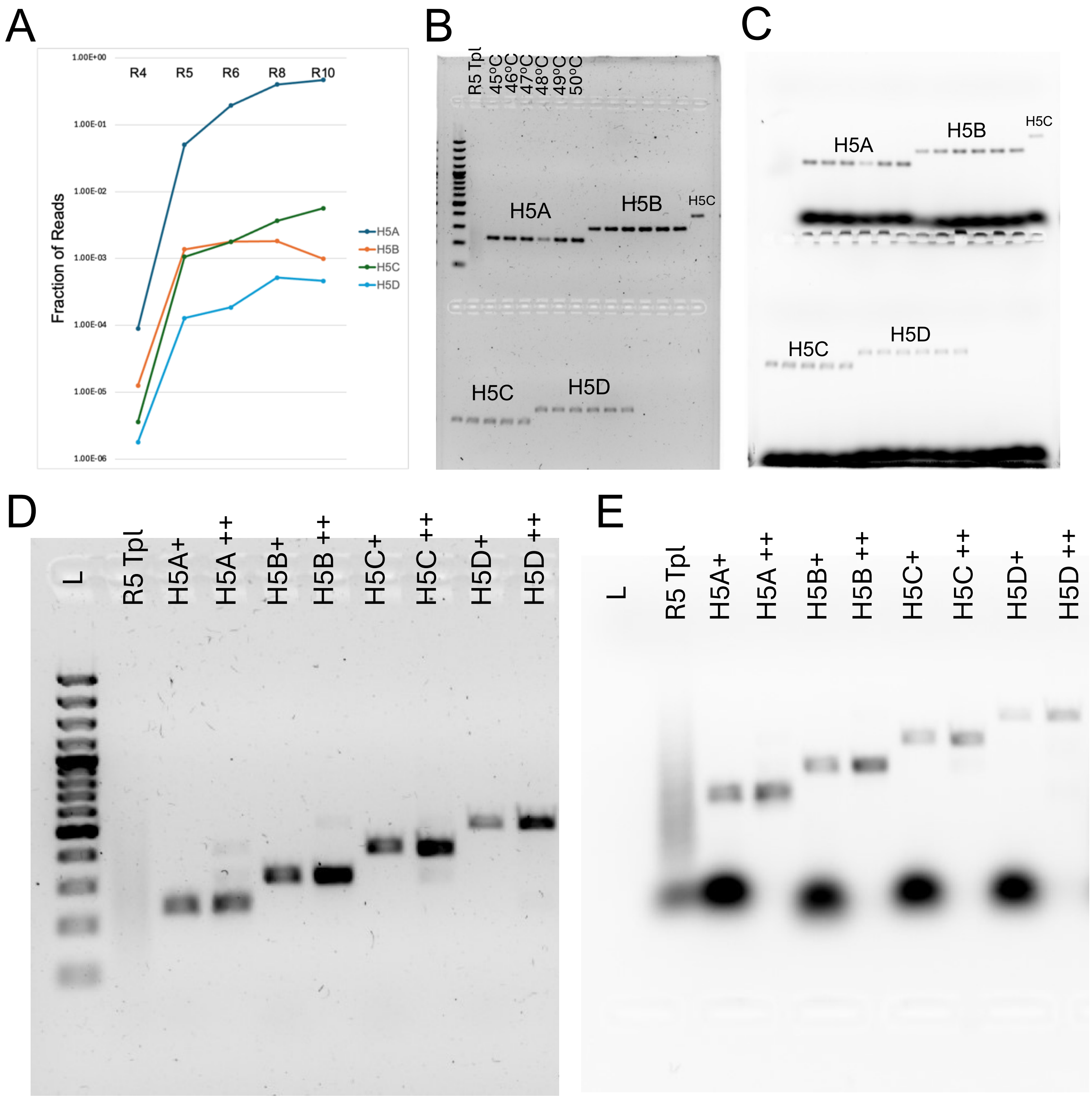


**Supplementary figure 4: Optimization of candidates selected from HEK293T nanopore sequencing of R-5.** (A)Enrichment profile of selected candidates H5A-H5D across successive selection rounds from R-4 to R-10 expressed as fraction of total sequencing reads. (B) SYBR Safe–stained 2% agarose gel showing amplification efficiency of selected candidates across temperature gradient. (C) Cy5 channel imaging of gel in panel-B to confirm incorporation of the Cy5-labeled primer. (D) SYBR Safe–stained 2% agarose gel showing amplification of candidates at optimized annealing temperature and their subsequent purification. (E) Cy5 channel imaging of gel in panel-D confirming removal of unincorporated primers after purification. *Abbreviations: Tpl is Template only, (+): PCR Amplification and (++): PCR amplification followed by Ampure purification*

A

| QUERY | SCORE | START | END | QSIZE | IDENTITY | CHROM | STRAND | START | END | SPAN |
|---|---|---|---|---|---|---|---|---|---|---|
| H5A | 165 | 1 | 184 | 208 | 95.2% | chr18 | - | 68074072 | 68074257 | 186 |
| H5A | 80 | 89 | 184 | 208 | 92.8% | chr9 | - | 129984888 | 129984987 | 100 |
| H5A | 78 | 90 | 184 | 208 | 91.6% | chr7 | - | 157064855 | 157064951 | 97 |
| H5A | 77 | 92 | 184 | 208 | 94.4% | chrX | - | 13176539 | 13176634 | 96 |
| H5A | 77 | 90 | 184 | 208 | 91.6% | chr8 | - | 129618282 | 129618380 | 99 |
| H5B | 230 | 2 | 242 | 283 | 98.0% | chrX | - | 116243695 | 116243936 | 242 |
| H5B | 199 | 2 | 242 | 283 | 92.4% | chr7 | - | 81034299 | 81034540 | 242 |
| H5B | 198 | 2 | 241 | 283 | 92.4% | chr14 | - | 36342126 | 36342366 | 241 |
| H5B | 198 | 3 | 246 | 283 | 91.0% | chr2 | + | 77657431 | 77815648 | 158218 |
| H5B | 197 | 2 | 246 | 283 | 90.3% | chr11 | - | 80139323 | 80139566 | 244 |
| H5C | 325 | 25 | 373 | 418 | 96.9% | chrUn_GL000224v1 | + | 822 | 1175 | 354 |
| H5C | 325 | 25 | 373 | 418 | 96.9% | chrUn_GL000224v1 | + | 7118 | 7471 | 354 |
| H5C | 320 | 25 | 373 | 418 | 97.7% | chrUn_KI270744v1 | + | 116180 | 116549 | 370 |
| H5C | 320 | 25 | 373 | 418 | 97.4% | chrUn_KI270744v1 | + | 122469 | 122830 | 362 |
| H5C | 320 | 25 | 373 | 418 | 97.4% | chrUn_GL000214v1 | + | 126232 | 126611 | 380 |
| H5D | 415 | 1 | 434 | 484 | 98.0% | chr11 | - | 24738612 | 24739045 | 434 |
| H5D | 25 | 27 | 51 | 484 | 100.0% | chr7 | - | 48618426 | 48618450 | 25 |
| H5D | 23 | 396 | 423 | 484 | 80.8% | chr10 | - | 89680776 | 89680801 | 26 |
| H5D | 23 | 249 | 273 | 484 | 96.0% | chr17 | + | 12077784 | 12077808 | 25 |
| H5D | 20 | 252 | 271 | 484 | 100.0% | chr20 | - | 11058500 | 11058519 | 20 |

B

**cDNA H5A**

```
TAAcGAAGAA GAAAAATAtc GacGGTGAGG CAGTAATTGC ACTAAATAAC  50
TATAGCAGAT TTCAGGGGGA GAGTTTAAAa CATAGATGTT GTATTAGTCC  100
ATTTTAACGC TGCTGATAAA GACATgTGGG AGACTGGGAA GAAAaAGGTT  150
TAATTTCACT TACAGcTCCA CATGGCTGGA GGCCgcccat cgtatctatt  200
atcgcttg
```

**cDNA H5B**

```
gGACTTCAGG AGTGGAACCa CAGACCTTCA CAGTGAGTGT TACAGCTCTT  50
AAAGGTGGCG CGTCCGGAGT TGTTTGTTCC TCTCGGTGGG TTtGTGGTCT  100
CACTGACTTC AGGAATGgAG TCTAGACCCT CGCAGTGAGT GTTACAGCTC  150
ATAAAAGTAG TGTGGACCCA AAGAGTGAcC AtCAGCAAGA TTTATTGTGA  200
AGAGCGAAAG AACAAAGCTT CCACAGTATG GAAGGGGACC CAcagctagc  250
ggatagactc caggacgcac tcaccaattg gcg
```

**cDNA H5C**

```
ccagcacgtg atatgtatat gtatCTCCCA GaGGCCCCAC TAATGTTCTA  50
ACTCGCAGAA GGAgGGAGTG TGTGTGCGTG TGTaCGTGTG CGTGTGTGTG  100
TGTGTGTTTG TGTtTGTGaG GTGTGAGGTA TGtGCCCCTT AAGaAAATGG  150
AAATCAACCA ACCAATGGGA aAGAgAGACA GACAGACAGA CAGAGATTCA  200
CTTGCCCAAc TGTTCTGTCC TGTCCTCTGA ATCCGCTTcC AAGTCGCAAG  250
ACGCTGTGAG CTCCAAGTCC ACGCAGAGTC CGCCAAACGC TCCGGCCGCT  300
GATCCGCTCC GCGAAGATCT GAGTACAGGC CAGCCAGGGT GGGTTTAAAT  350
AGCCTCGGGC GCAGCCTAGC AGCccatacg tatctagatg ccaatatcta  400
tacatgtacg cgtgtagg
```

**cDNA H5D**

```
TGACCTTACC ATACCTCATA TGTTCAATTA ACACACTATA TTTTAAAACA  50
GACTCAGATC TTGAAAACAA GATATAAGGT CCAATCCCTA TACATGGTGC  100
GTAAGGTGGC TTTGGGGAAT ACAGTAGAAA CCTAGTTTTC AGCATGGCTC  150
TCAGaAAACA CCATCCCTCC TGCCTGTGGT AAgTATTCGT AGTAAGAGCT  200
GAAACGGTGG GGTCAAGaGA ACATCAcTTC GGGTGAGAAA GTTTGTTTCA  250
TACTTTATTT CTCACAATTC TcGCcGTCAa GAGAGGAGTT TGATTCATCA  300
ACTTTGGGAT GCATTAGAAT AACCCGTGGG ATGGATTAAA TAGATTCCAT  350
GGCCCTGCTC CTGAGAGATC CTTATTCAGT GGATGTGacG TGCGACTTAA  400
CTATCACTTA GCTTAATTTA CTTCACAGTC AGTGtacata tgcacactgc  450
tgtgtataga actccatacg tatctactca ctcc
```

**Supplementary figure 5: Genome mapping and alignment analysis of selected structural candidates.** (A) BLAT analysis summary displaying the top five alignment hits for candidates H5A–H5D against the human reference genome (GRCh38). Higher scores indicate greater sequence homology between the candidate query and the host genome. (B) Detailed alignment maps of the primary hit for each candidate. The matching bases between the query and genomic sequences are indicated by capitalized blue letters. Light blue highlights mark the boundaries of gaps or transitions (such as potential splice sites) within the sequence alignment and black non capitalized letters marks no significant alignments and is mostly on the 3 ' end of the query sequence.

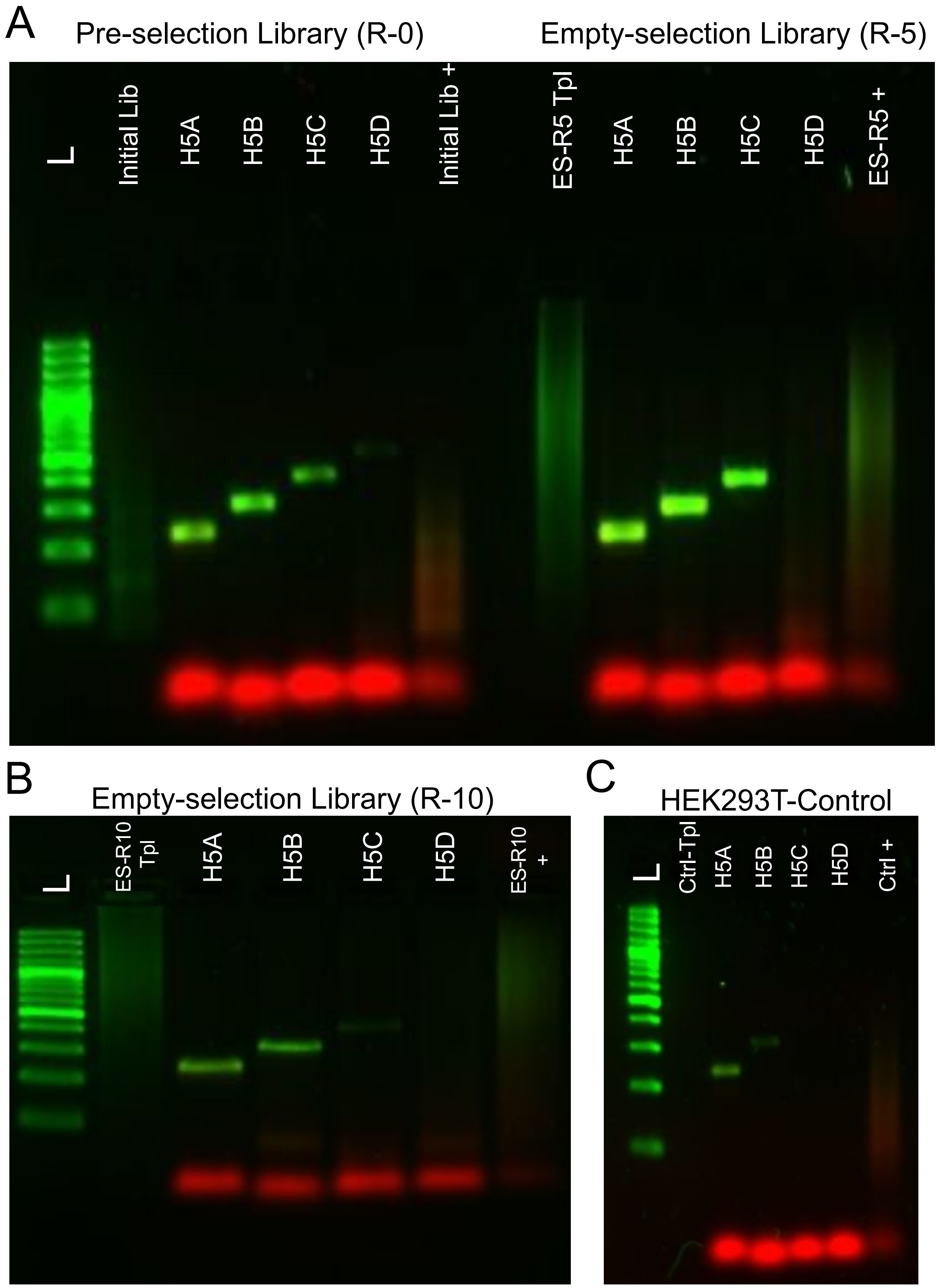


**Supplementary figure 6:** ***Verification of genomic origin in control samples.*** Agarose gels stained with SYBR Safe were imaged using the blot setting of a gel documentation system, with AF488 displayed in green and Cy5 in red. (A) Initial library (pre-selection library) showing all candidates with a faint H5D band, and empty selection R-5, representing the initial library subjected to multiple rounds of amplification and AMPure purification without cell stimulation. (B) Empty selection R-10 showing faint bands for H5A and H5B, with no detectable H5C and H5D. (C) Amplification from non-treated HEK control cells to assess cellular contamination, showing a band only for H5A and very faint H5B. *Abbreviations: Initial Lib: initial library before any cell stimulation, Initial Lib+: PCR amplified initial lib using amplification loop Cy5 primers, ES-R5 Tpl: Empty selection library after five rounds of amplification and purification used as template, ES-R10: Empty selection library after ten round of*

*amplification and purification used as template, Ctrl Tpl: HEK293T non-treated control purified lysate used as template.*

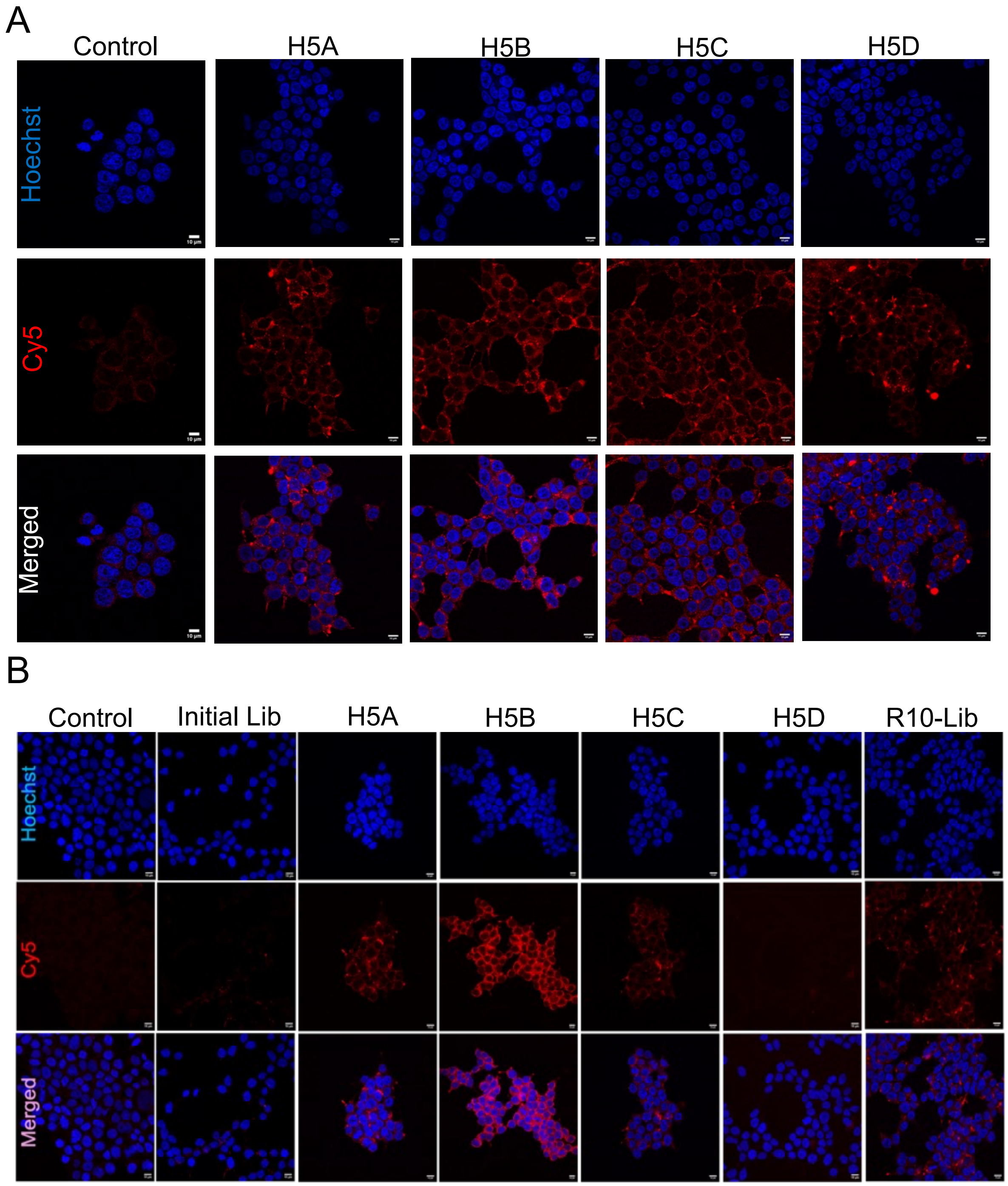


**Supplementary figure 7: Cellular uptake of selected candidates in HEK293T cells.** (A) representative laser scanning confocal images of fixed cells stimulated with 10 nM of individual candidates amplified from the Round-5 template. (B) Cellular uptake of candidates amplified from Round-10 template and control samples. Preselection library (initial lib) was the library used for the initial round of cell stimulation and Round-10 library (R10-Lib) was the library after ten rounds of selection were controls amplified using Cy5-labeled UMI primers. Nuclei are stained with Hoechst (blue), and Cy5-labeled DNA candidates are shown in red; the merged panel displays combined blue and red channels. Scale bar: 10 µm

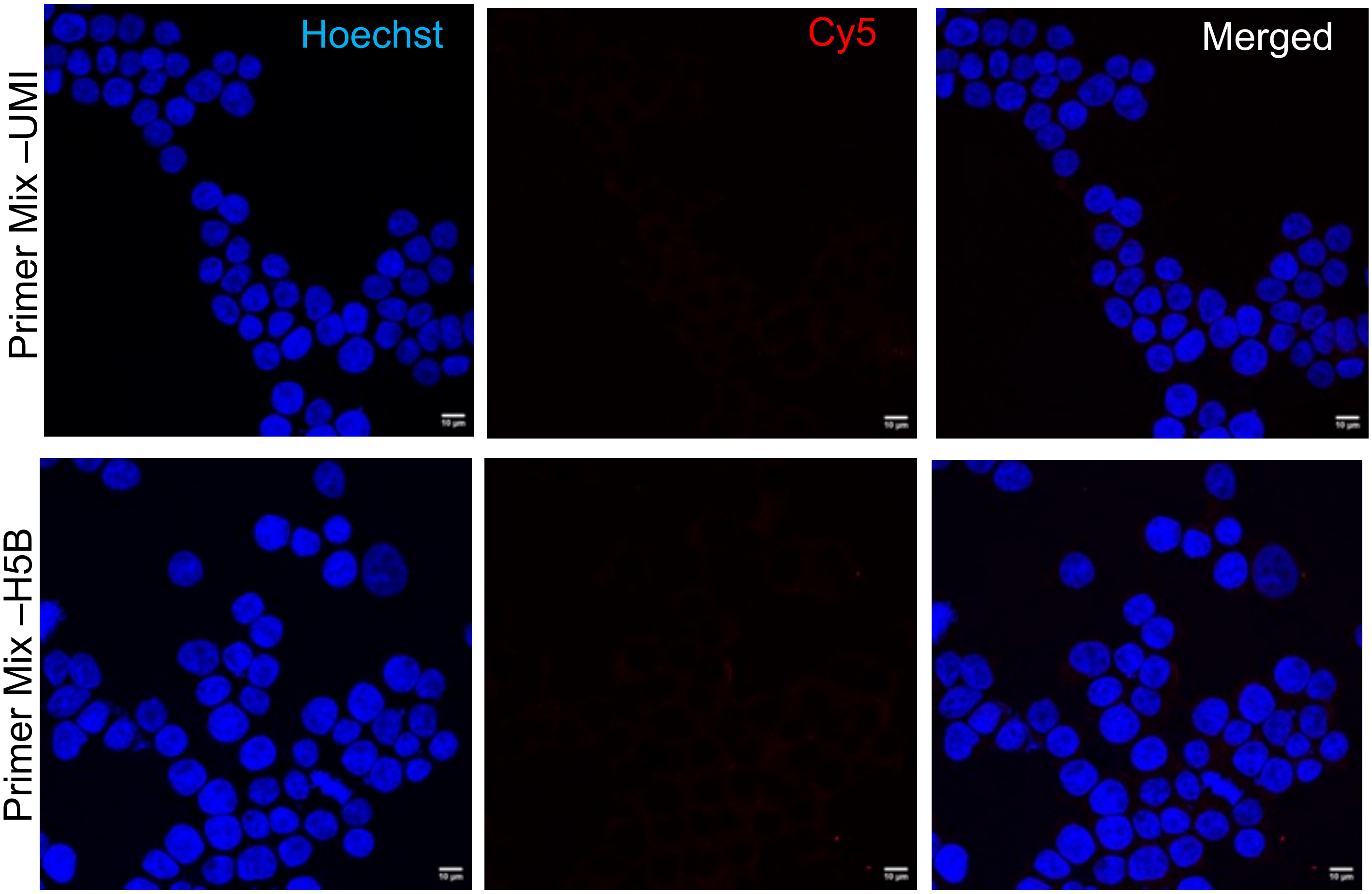


**Supplementary figure 8: Assessment of potential false-positive signals from Cy5-labeled primer mixtures.** Confocal microscopy images were acquired to monitor the cellular uptake of residual primers from UMI and H5B sets. Primers were pooled and added to cells at a final concentration of 50 nM to evaluate whether unreacted primer mixtures contribute to false-positive intracellular signals. The red channel represents the Cy5 signal, the blue channel represents Hoechst-stained nuclei, and the merged panel shows the overlay of both channels. Scale bar: 20 μm.

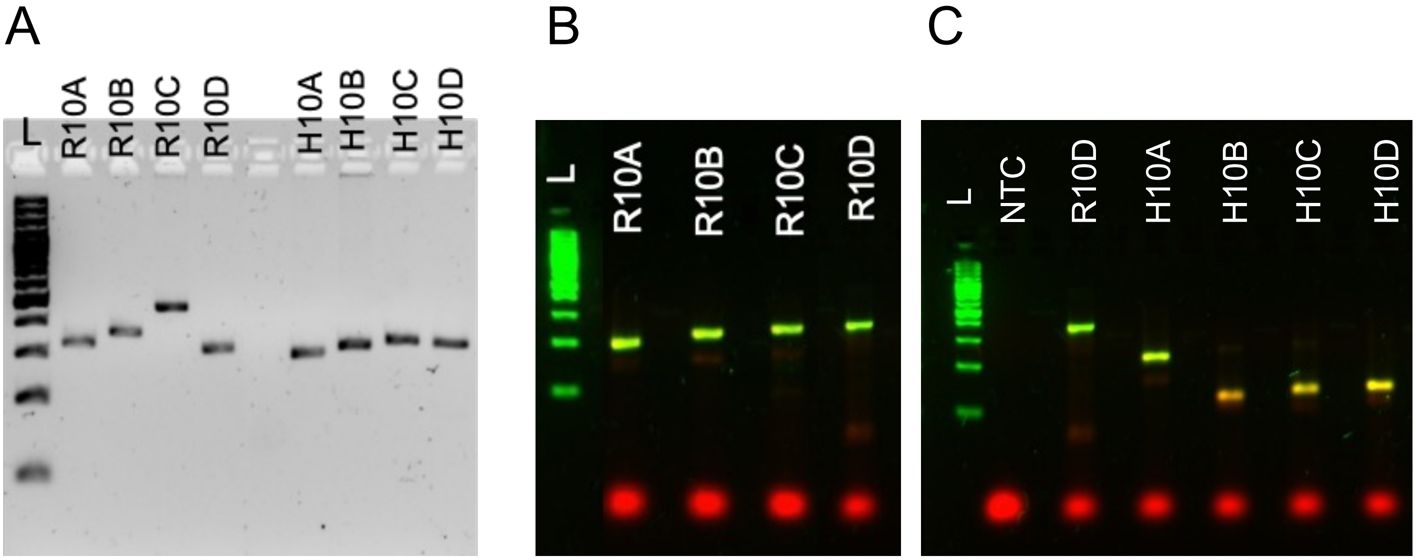


**Supplementary figure 9: Optimization of HEK293T and RAW264.7 selected candidates from R-10 Illumina data**. (A) 2% agarose gel stained with SYBR safe demonstrating the purity of the constructs that will be used as template for subsequent amplification. (B) Amplification of RAW264.7 selected candidates. Agarose gel stained with SYBR safe and imaged in blot mode; with green color indicates the SYBR stained DNA and red color indicates Cy5 primer incorporation. (C) Amplification of HEK293T selected candidates under same conditions as in (B). NTC is no template control and showed no detectable amplification. All the candidates showed successful amplification without any non-specific amplification.

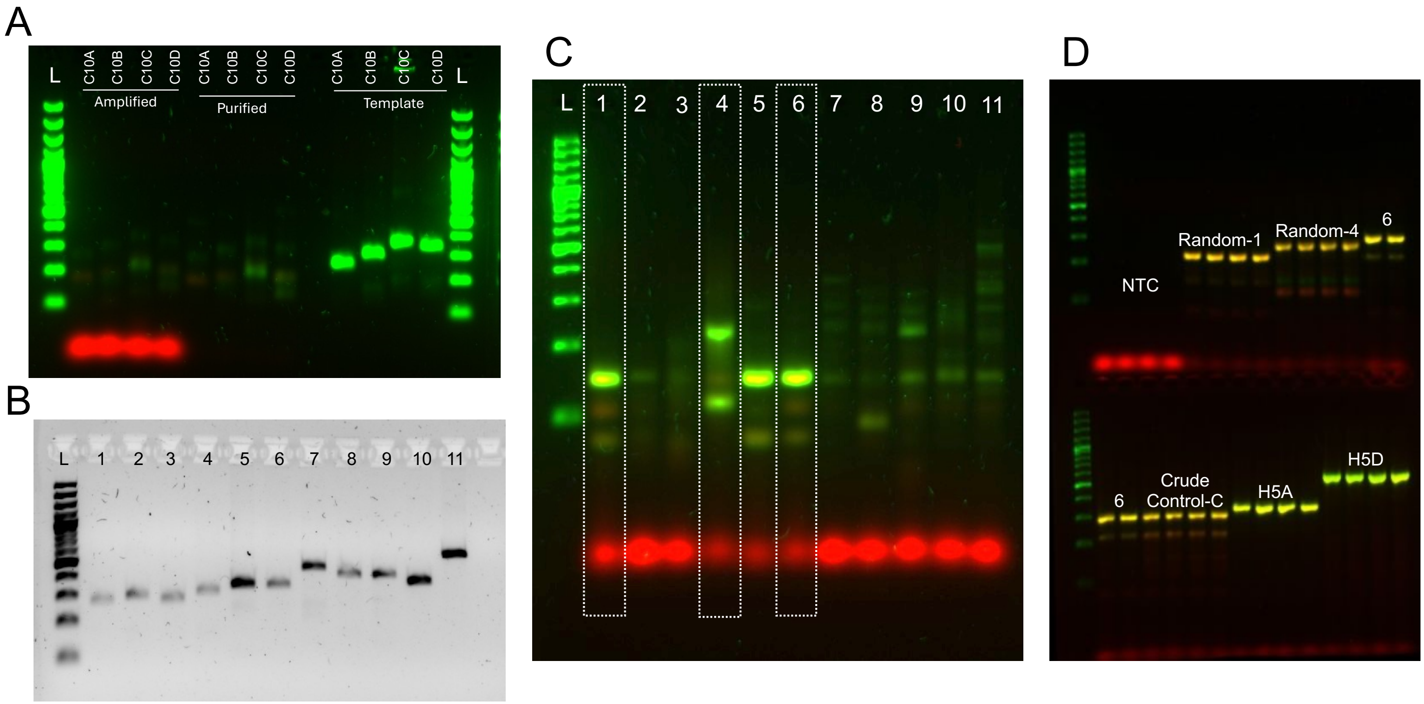


**Supplementary figure 10: Optimization of control candidates for cellular study. (A)** Agarose gel electrophoresis of crude control candidates (C10 A-D) visualized using SYBR Safe (green; AF488 channel) and Cy5 fluorescence (red). Lane designations indicate: "Amplified" (crude PCR product), "Purified" (samples after Ampure bead purification showing removal of residual primers), and "Template" (stock template loaded to assess initial heterogeneity). **(B)** SYBR Safe-stained agarose gel of newly selected random candidates (Lanes 1–9) and control candidates H5A (Lane 10) and H5D (Lane 11). **(C)** PCR amplification of random candidates from template shown in panel B using Cy5-labeled primers, visualized via SYBR Safe (green; AF488) and Cy5 (red) channels. Highlighted lanes indicate the specific candidates selected for further optimization and cellular study. **(D)** Temperature optimization of selected random control candidates; NTC denotes the No Template Control.

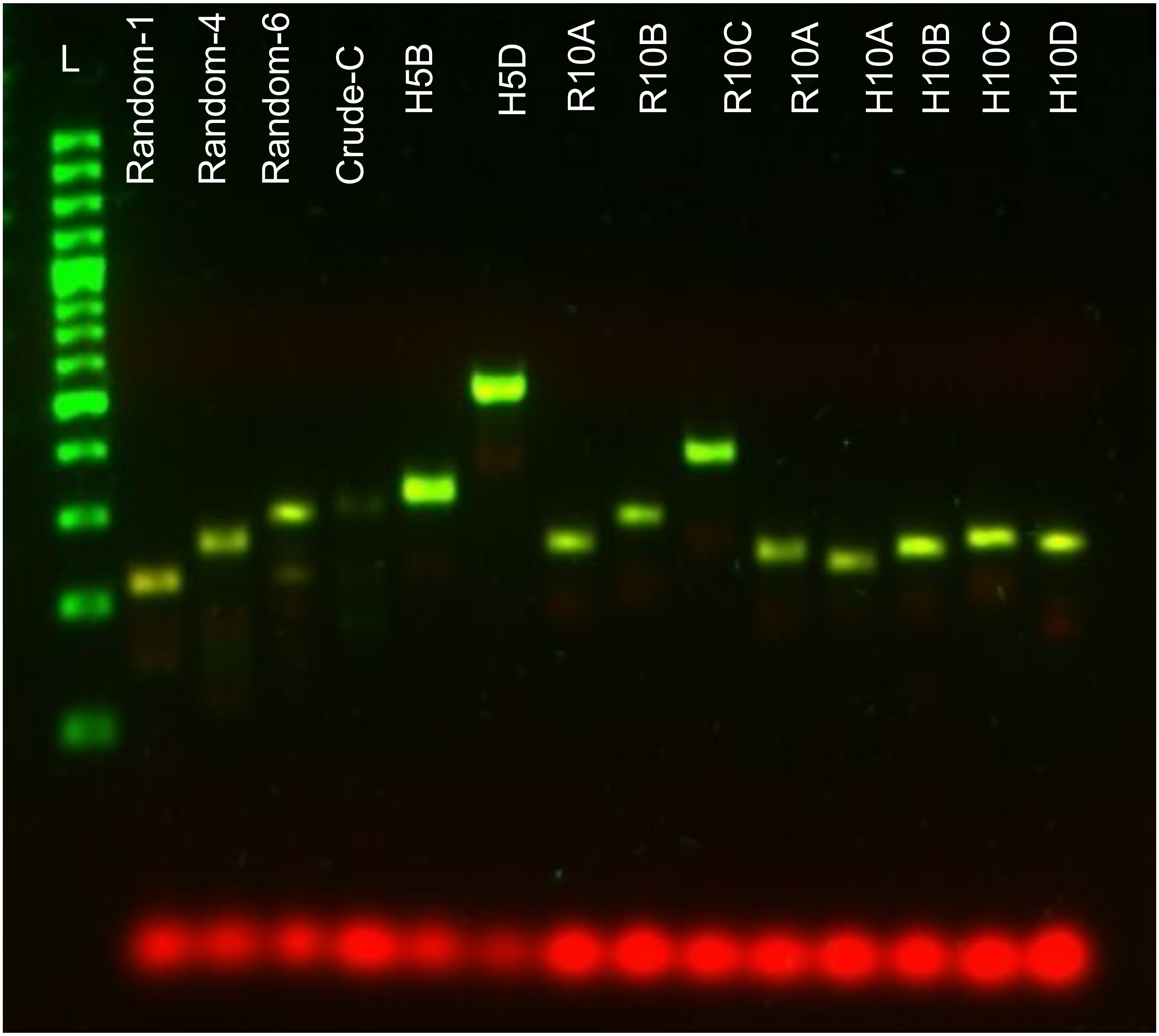


**Supplementary figure 11: Agarose gel verification of finalized candidates.** Amplified candidates were resolved on an agarose gel to verify assembly and purity. Samples were loaded as indicated on the respective lanes and electrophoresed for 45 minutes at 100V. Lane L denotes the 100 bp DNA ladder. The green signal represents SYBR Safe DNA stain acquired under the AF488 channel, while the red signal represents Cy5 detection.

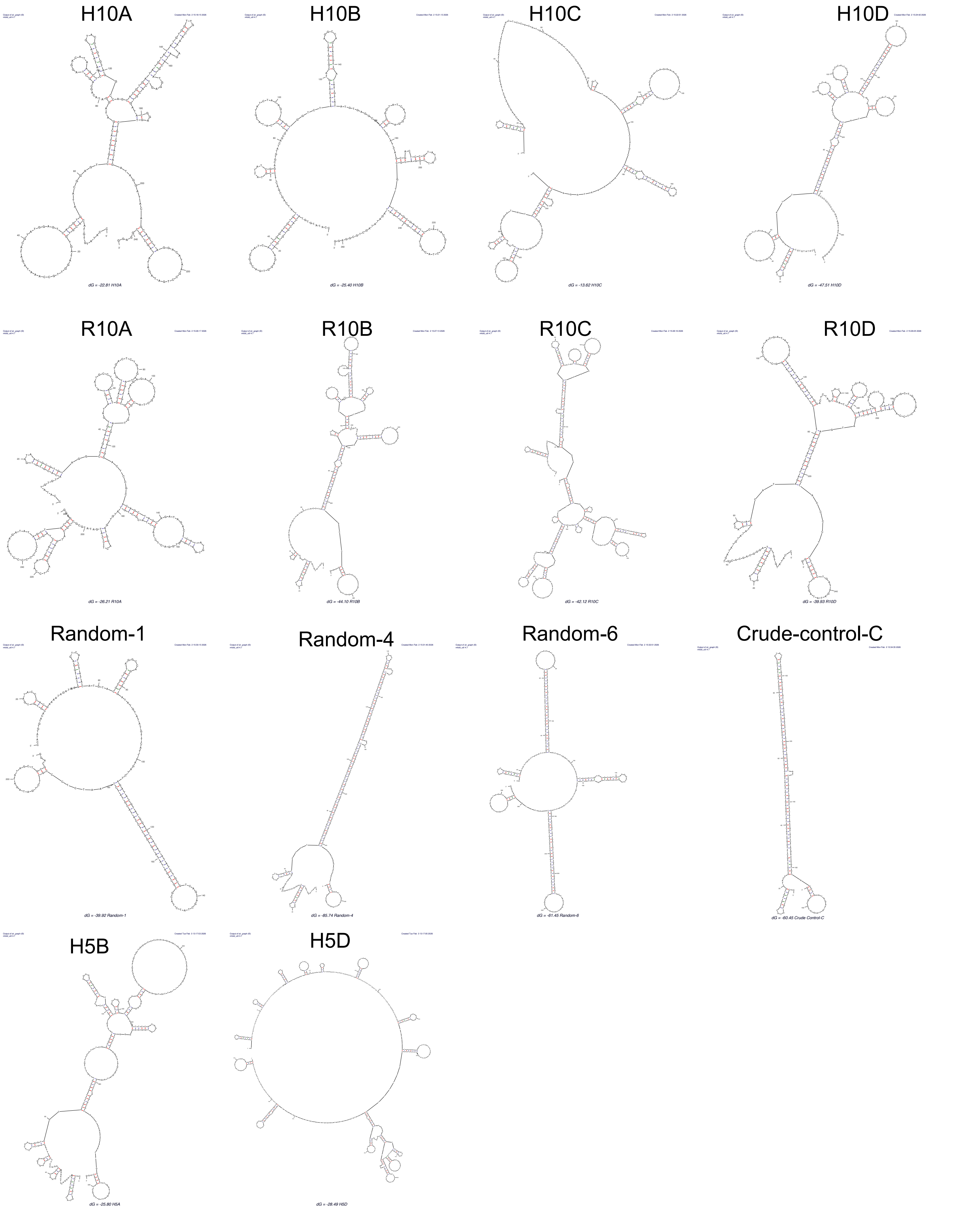


**Supplementary figure 12:** In Silico Folding Analysis and Structural Characterization of Finalized DNA Sequences predicted by m-Fold

| Sequence Name | Length (nt) | Energy (ΔG) | Tracebacks | Helix Count | Stability Density (ΔG/nt) |
|---|---|---|---|---|---|
| Random-4 | 254 | -85.7 | 1 | 107 | 0.337 |
| Random-6 | 288 | -61.5 | 3 | 116 | 0.214 |
| Crude-Control-C | 233 | -61.5 | 1 | 37 | 0.331 |
| H10D | 271 | -47.5 | 2 | 104 | 0.175 |
| R10B | 296 | -44.1 | 2 | 60 | 0.149 |
| R10D | 256 | -39.9 | 1 | 74 | 0.156 |
| Random-1 | 213 | -39.9 | 2 | 49 | 0.187 |
| H10A | 244 | -21.0 | 3+ | 38 | 0.086 |
| H5D | 528 | -28.5 | 1 | 116 | 0.054 |
| R10C | 395 | -42.1 | 3 | 152 | 0.107 |
| R10A | 263 | -26.2 | 2 | 41 | 0.099 |
| H10B | 262 | -25.4 | 3 | 58 | 0.090 |
| H5B | 327 | -25.8 | 2 | 44 | 0.079 |
| H10C | 279 | -20.8 | 3 | 51 | 0.075 |

**Supplementary Table 5:** Structural characteristics of finalized candidates based on mfold secondary structure predictions

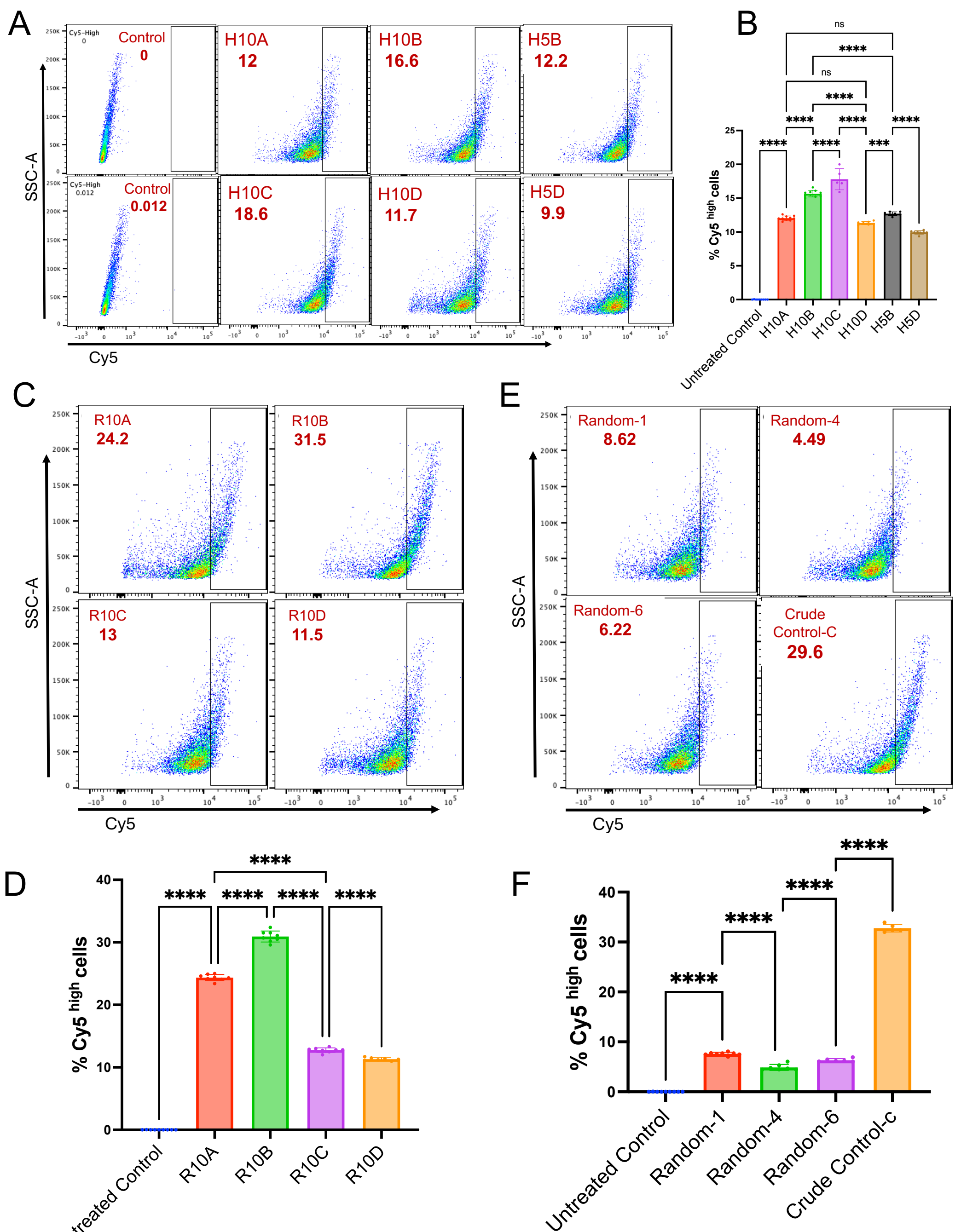


**Supplementary figure 13: Quantification of DNA nanostructure internalization via flow cytometry in RAW264.7 cells.** Data shown represent the gated Cy5-high cell population. **(A)** Representative flow cytometry scatter plots of RAW264.7 cells treated with DNA nanostructure candidates selected from HEK293T screens. **(B)** Quantification of Cy+ cells calculated from scatter plots shown in (A). **(C)** Representative flow cytometry scatter plots of cells treated with candidates selected from RAW264.7-specific screens. **(D)** Quantification of Cy+ cells calculated from scatter plots shown in (C). **(E)**Representative flow cytometry scatter plots of cells treated with randomly selected candidates. **(F)**Quantification of Cy+ cells calculated from scatter plots shown in (E). For all flow cytometry experiments, 10,000 events were recorded per sample. Samples were acquired in quadruplets, and data represent two independent experiments (N = 2). Error bars indicate mean ± SD. Statistical significance was determined by ordinary one-way ANOVA (****P < 0.0001, ns: non-significant).

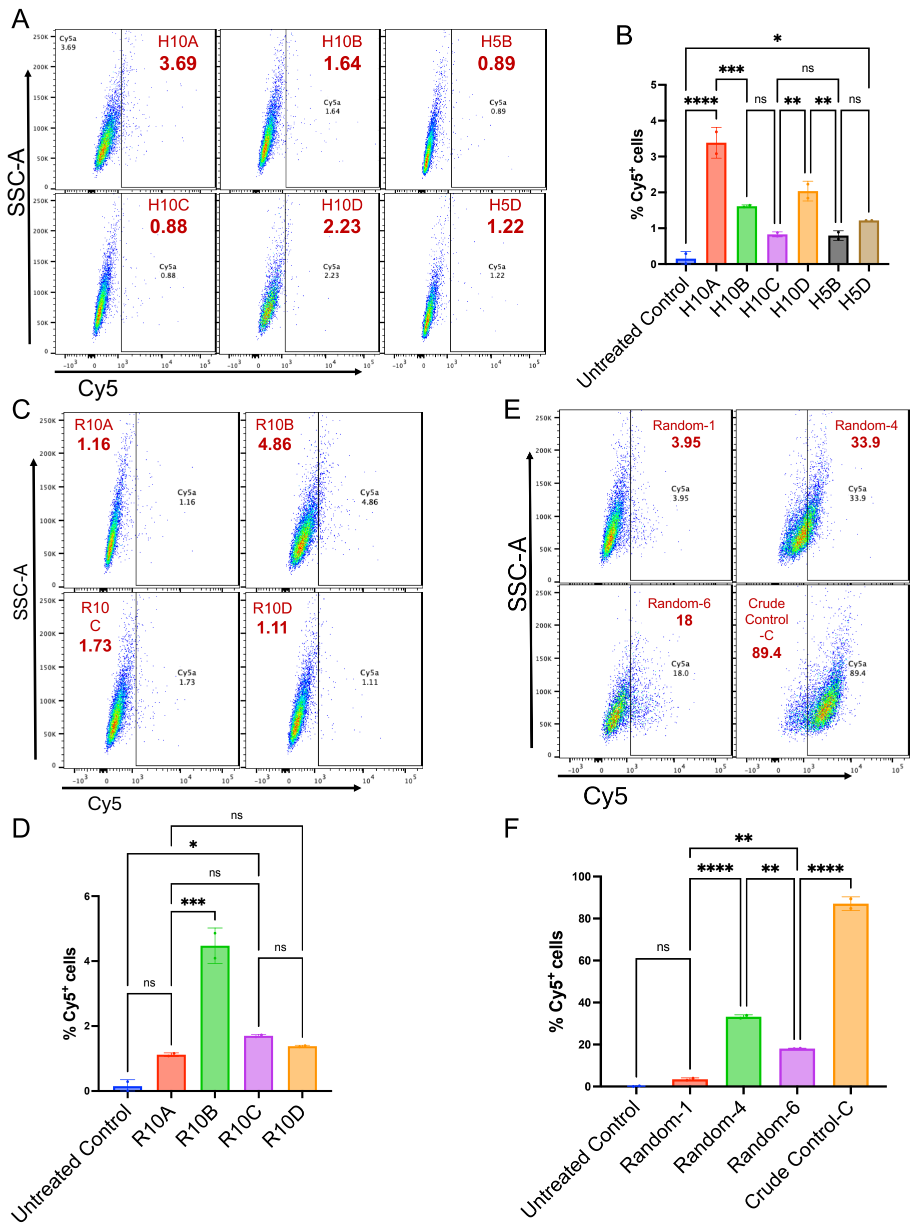


**Supplementary figure 14: Quantification of DNA nanostructure internalization via flow cytometry in A549 cells. (A)** Representative flow cytometry scatter plots of A549 cells treated with DNA nanostructure candidates selected from HEK293T screens. **(B)** Quantification of Cy+ cells calculated from scatter plots shown in (A). **(C)** Representative flow cytometry scatter plots with candidates selected from RAW264.7-specific screens. **(D)** Quantification of Cy+ cells calculated from scatter plots shown in (C). **(E)** Representative flow cytometry scatter plots of cells treated with randomly selected candidates. **(F)** Quantification of Cy+ cells calculated from scatter plots shown in (E). For all flow cytometry experiments, 10,000 events were recorded per sample. Samples were acquired in duplicate, and data represent two independent experiments (N = 2). Error bars indicate mean ± SD. Statistical significance was determined by ordinary one-way ANOVA (****P < 0.0001,***P

< 0.0005, **P < 0.0012, ns: non-significant).

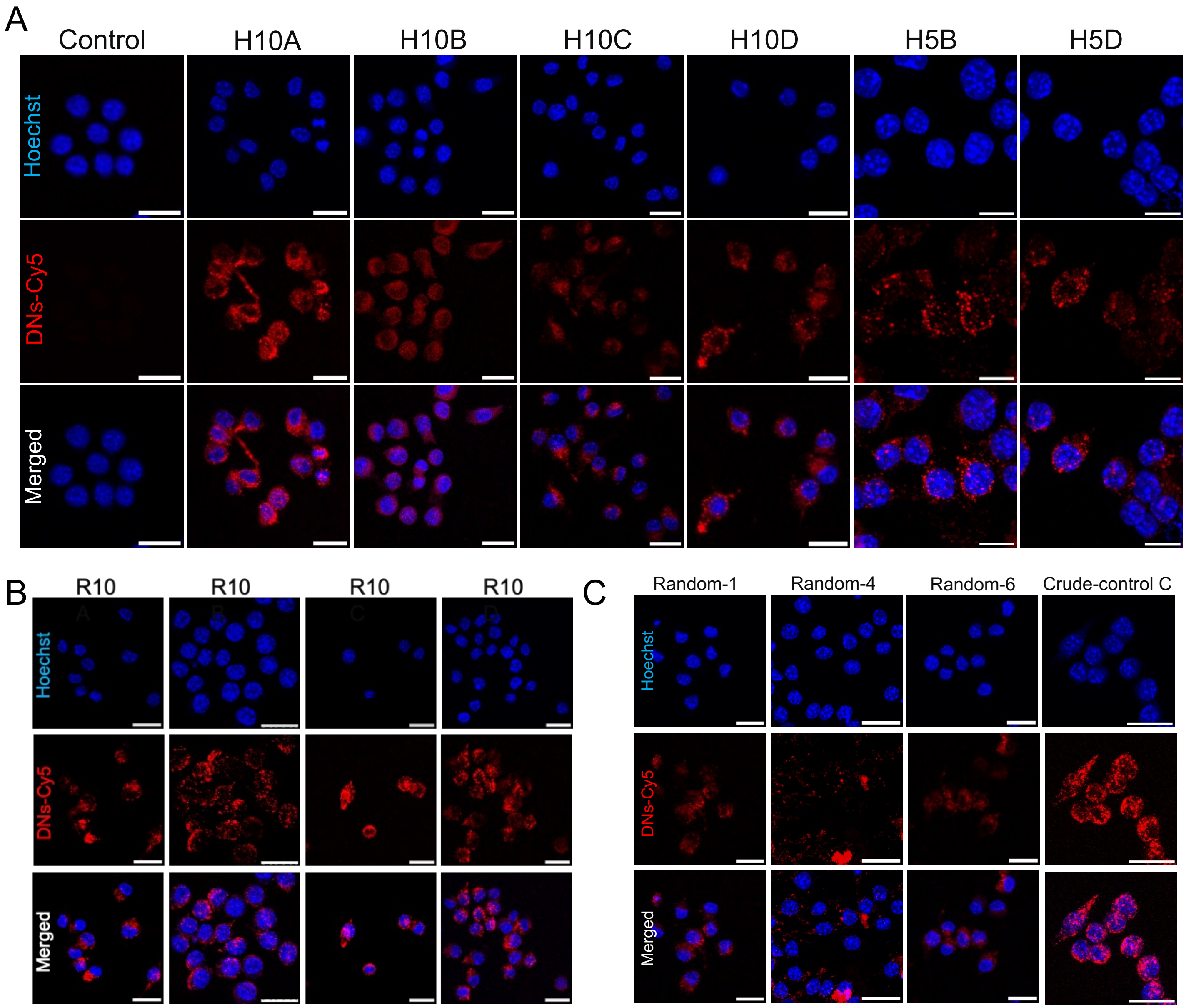


**Supplementary figure 15: Qualitative assessment of cellular uptake of final candidates in RAW264.7 using laser scanning confocal microscopy. (A)** Confocal images of cells treated with HEK293T-selected DNA nanostructure (DN-Cy5). **(B)** Confocal images of cells treated with RAW cell-selected DNA nanostructure. **(C)** Confocal images of RAW264.7 cells treated with randomly selected DNA nanostructure. For all panels, the red channel represents DN-Cy5 uptake, and the bottom panels represent merged images of all channels with nuclei stained with Hoechst dye (blue). Scale bar: 20 µm.

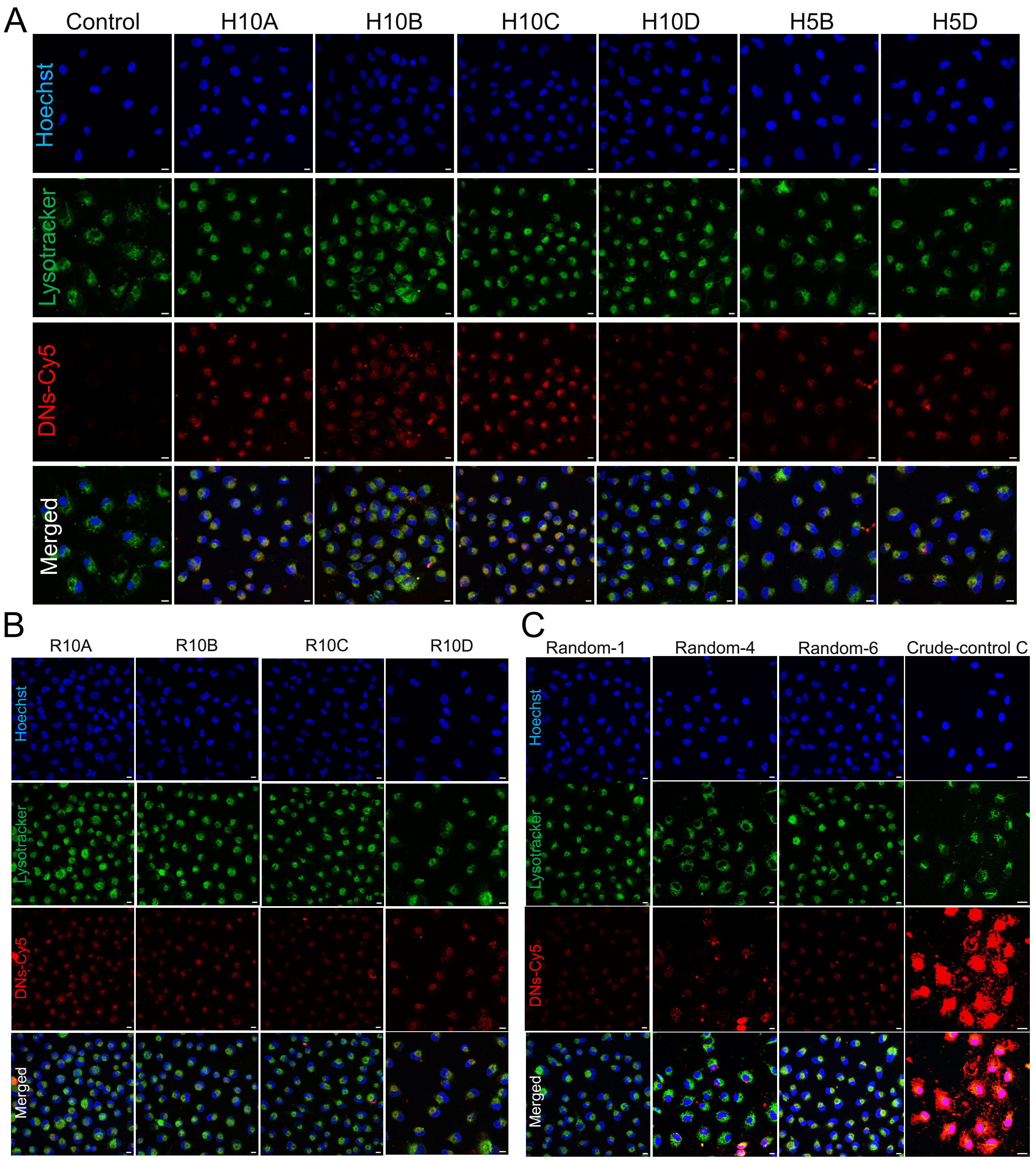


**Supplementary figure 16: Qualitative assessment of cellular uptake of final candidates in A549 cells using confocal microscopy. (A)** Confocal images of A549 cells treated with HEK293T-selected DNA nanostructure (DN-Cy5). **(B)** Confocal images of RAW cell-selected candidates. **(C)** Confocal images of randomly selected candidates. For all panels, the red channel represents DN-Cy5 uptake, the green channel indicates LysoTracker Green staining for lysosomes, and the bottom panels represent merged images of all channels with nuclei stained with Hoechst dye (blue). Scale bar: 15 μm.
